# Dominant role of adult neurogenesis-induced structural heterogeneities in driving plasticity heterogeneity in dentate gyrus granule cells

**DOI:** 10.1101/2021.08.22.457267

**Authors:** Sameera Shridhar, Poonam Mishra, Rishikesh Narayanan

## Abstract

Neurons and synapses manifest pronounced variability in the amount of plasticity induced by identical activity patterns. The mechanisms underlying such plasticity heterogeneity, implicated in context-specific resource allocation during encoding, have remained unexplored. Here, we employed a systematic, unbiased, and physiologically constrained search to identify the mechanisms behind plasticity heterogeneity in dentate gyrus granule cells. We found that each of intrinsic, synaptic, and structural heterogeneities independently yielded heterogeneous plasticity profiles obtained with two different induction protocols. However, prior predictions about strong relationships between neuronal intrinsic excitability and plasticity emerged only when adult-neurogenesis-induced structural heterogeneities were accounted for. Strikingly, despite the concomitant expression of heterogeneities in structural, synaptic, and intrinsic neuronal properties, similar plasticity profiles were attainable through synergistic interactions among these heterogeneities. Importantly, consequent to strong relationships with intrinsic excitability measurements, we found that synaptic plasticity in the physiological range was achieved in immature cells despite their electrophysiologically-observed weak synaptic strengths. Together, our analyses unveil the dominance of neurogenesis-induced structural heterogeneities in driving plasticity heterogeneity in granule cells. Broadly, these analyses emphasize that the mechanistic origins of and the implications for plasticity heterogeneities need quantitative characterization across brain regions, particularly focusing on context-specific encoding of learned behavior.

## INTRODUCTION

Neurons and synapses of the same subtype receiving identical plasticity-inducing activity patterns do not manifest identical levels of plasticity, instead exhibiting pronounced *plasticity heterogeneity* across synapses and neurons. There are several lines of evidence from *in vitro* and *in vivo* electrophysiological experiments for such plasticity heterogeneity, spanning different neuronal and synaptic subtypes (Bliss and Lomo, 1973; Greenstein et al., 1988; Pavlides et al., 1988; Shors and Dryver, 1994; Wang et al., 1997; Beck et al., 2000; Davis et al., 2004; McHugh et al., 2007; Koranda et al., 2008; Sjostrom et al., 2008; Kobayashi et al., 2013; Larson and Munkacsy, 2015; Li et al., 2017; Rathour and Narayanan, 2019). Although such plasticity heterogeneity has typically been overlooked in analyzing the impact of plasticity protocols, a growing body of experimental evidence has identified crucial roles for plasticity heterogeneity in neural encoding and storage. Specifically, the ability of neurons and synapses to undergo differential plasticity is critical for *context-specific recruitment/allocation of a subset of* neurons and synapses during encoding processes (Schmidt-Hieber et al., 2004; Ge et al., 2007; Dieni et al., 2013; Aimone et al., 2014; Yiu et al., 2014; Park et al., 2016; Josselyn and Frankland, 2018; Lodge and Bischofberger, 2019; Pignatelli et al., 2019; Josselyn and Tonegawa, 2020; Lau et al., 2020; Huckleberry and Shansky, 2021). The lack of plasticity heterogeneity would result in a scenario where all neurons and synapses undergo similar amount of plasticity for any given context, thus erasing the possibility of sparse and context-specific recruitment of neural resources. Despite these well-recognized roles of plasticity heterogeneity in context-specific resource allocation, the *mechanisms* underlying these heterogeneities have not been assessed. Furthermore, although there are postulates and lines of evidence for heterogeneities in intrinsic excitability playing a role in determining selective resource allocation (Schmidt-Hieber et al., 2004; Ge et al., 2007; Dieni et al., 2013; Aimone et al., 2014; Yiu et al., 2014; Park et al., 2016; Josselyn and Frankland, 2018; Lodge and Bischofberger, 2019; Pignatelli et al., 2019; Josselyn and Tonegawa, 2020; Lau et al., 2020; Huckleberry and Shansky, 2021), the quantitative link between such cellular-scale heterogeneities and plasticity heterogeneity has not been assessed.

Granule cells (GC) in the dentate gyrus (DG) offer an efficient system for addressing questions on the cellular mechanisms underlying plasticity heterogeneity. First, the pronounced biophysical heterogeneities in these cell types have been electrophysiologically well-characterized (Schmidt-Hieber et al., 2004; Overstreet-Wadiche et al., 2006b; Pedroni et al., 2014; Heigele et al., 2016; Mishra and Narayanan, 2020a). Second, plasticity experiments involving granule cells have revealed the manifestation of heterogeneities in the amount of synaptic plasticity induced for the same activity protocols (Bliss and Gardner-Medwin, 1973; Bliss and Lomo, 1973; Greenstein et al., 1988; Pavlides et al., 1988; Shors and Dryver, 1994; Wang et al., 1997; Beck et al., 2000; Davis et al., 2004; McHugh et al., 2007; Koranda et al., 2008; Kobayashi et al., 2013; Larson and Munkacsy, 2015). Third, these intrinsic and plasticity heterogeneities are further amplified by the expression of adult neurogenesis, resulting in immature neurons that manifest increased excitability, reduced synaptic connectivity, lesser dendritic arborization, and hyperplasticity that translates to lower threshold for plasticity induction (Schmidt-Hieber et al., 2004; Ge et al., 2007; Dieni et al., 2013; Aimone et al., 2014; Li et al., 2017; Lodge and Bischofberger, 2019; Huckleberry and Shansky, 2021). Finally, there are lines of evidence for a critical role of plasticity heterogeneity in engram formation, response decorrelation, and resource allocation in the dentate gyrus, with postulates about the role of intrinsic excitability in governing plasticity rules and selective resource allocation (Aimone et al., 2014; Yiu et al., 2014; Park et al., 2016; Josselyn and Frankland, 2018; Lodge and Bischofberger, 2019; Mishra and Narayanan, 2019; Pignatelli et al., 2019; Josselyn and Tonegawa, 2020; Lau et al., 2020; Huckleberry and Shansky, 2021). Together, granule cells allowed us to place plasticity heterogeneity within a strong functionally relevant context of engram formation and response decorrelation, providing an efficient substrate for assessing the impact of well-characterized biophysical and structural heterogeneities on the emergence of plasticity heterogeneity.

In this study, we systematically explored the cellular-scale origins of heterogeneities in the synaptic plasticity profiles of DG granule cells through an unbiased exploration of heterogeneities in their intrinsic, synaptic, and structural properties. We ensured that our analyses associated with each of these heterogeneities were constrained by characteristic physiological properties of granule cells spanning the immature-to-mature GC continuum. We assessed the impact of these forms of heterogeneities on plasticity profiles obtained with two well-established protocols for inducing synaptic plasticity: the 900-pulses protocol spanning a range of induction frequencies (Wang et al., 1997; Koranda et al., 2008; Kobayashi et al., 2013), and the theta-burst stimulation protocol (Greenstein et al., 1988; Pavlides et al., 1988; Shors and Dryver, 1994; Beck et al., 2000; Davis et al., 2004; McHugh et al., 2007; Larson and Munkacsy, 2015). We found that each form of intrinsic, synaptic, and structural heterogeneity independently resulted in plasticity heterogeneities, with either protocol for plasticity induction. Importantly, when immature and mature neuron populations were individually analyzed, we found that heterogeneities in intrinsic excitability were insufficient to impose strong constraints on plasticity-related measurements. However, when the entire population covering mature and immature cells were considered *together*, it was clear that there were strong relationships between intrinsic excitability and measurements associated with synaptic plasticity.

From the perspective of degeneracy, we show that the expression of heterogeneities in all of structural, synaptic, and intrinsic neuronal properties does not necessarily have to translate to heterogeneities in synaptic plasticity profiles. Specifically, we demonstrate that very *similar* plasticity profiles could be achieved with disparate combinations of neuronal passive properties, ion-channel properties, calcium-handling mechanisms, synaptic strength, and neural structure of DG granule cells of different ages. When observed independently, these properties manifested widespread heterogeneities with no pair-wise relationships, but when seen together, these heterogeneities synergistically interacted with each other to achieve the functional goal of degeneracy in synaptic plasticity profiles. These analyses extend degeneracy in DG granule cells to the *concomitant* emergence of plasticity profiles and of several neural intrinsic properties, while incorporating all three forms of cellular-scale heterogeneities spanning different age groups of GCs in a physiologically constrained manner. Strikingly, these analyses also showed that synaptic plasticity in the useful physiological range could be achieved in immature cells even with the weak synaptic strengths that they are known to express, owing to strong relationships with intrinsic excitability measurements.

Together, our analyses demonstrate that synergistic interactions between intrinsic, synaptic, and structural heterogeneities drive plasticity heterogeneity in dentate gyrus granule cells. Our results also highlighted the dominance of structural heterogeneities, introduced by adult neurogenesis, in introducing plasticity heterogeneity that is essential for context-specific resource allocation in the dentate gyrus. From a broader perspective, our analyses call for systematic characterization and analyses of plasticity heterogeneities across different brain regions, probing their mechanistic origins and their implications for context-specificity that is crucial for efficacious neural coding of learned behavior and memory storage.

## MATERIAL AND METHODS

The dentate gyrus GC population manifests significant amount of inherent intrinsic heterogeneity, which is further amplified by the expression of adult neurogenesis. Therefore, at any specified time, there are populations of neurons at different stages of maturation manifesting age-dependent structural heterogeneity. These adult-born granule cells at different stages of maturation exhibit heterogeneities in neuronal properties (intrinsic heterogeneity), in synaptic connections (synaptic heterogeneity), and structural properties including dendritic arborization and surface area (structural heterogeneity). In this study, our goal is to explore the impact of these heterogeneities on synaptic plasticity profiles, employing conductance-based models for DG granule cells. Assessment of plasticity profiles involve long-term simulations and the complexities associated with incorporating different forms of heterogeneities in *a population of conductance-based models* (as opposed to a single model with fixed structure and fixed synaptic strengths) implied large computational costs. Thus, we employed single-compartmental conductance-based models to assess the impact of different forms of biophysical and structural heterogeneities on synaptic plasticity induced through two extensively employed plasticity-induction protocols.

### Heterogeneities in intrinsic properties of a physiologically constrained granule cell model population

Granule cells in the dentate gyrus manifest pronounced heterogeneities in their intrinsic properties (Lubke et al., 1998; Aradi and Holmes, 1999; Santhakumar et al., 2005; Krueppel et al., 2011; Mishra and Narayanan, 2020a). The physiologically constrained conductance-based heterogeneous population of granule cell model was obtained from an earlier study (Mishra and Narayanan, 2019). The details of building this population of models that manifested characteristic electrophysiological properties of granule cells, employing a multi-parametric multi-objective stochastic search (MPMOSS) algorithm are identical to the previous study (Mishra and Narayanan, 2019). Briefly, the dimensions of single cylindrical base model were set to 63 μm diameter and 63 μm length, to achieve a passive input resistance of 305 MΩ, matching the experimental value of 309 ± 14 MΩ (Chen, 2004). The resting membrane potential of model cell was set to −75 mV, with specific membrane resistance (*R*_m_) of 38 kΩ.cm^2^ and specific membrane capacitance (*C*_m_) of 1 μF.cm^−2^. These passive parameters resulted in a charging time constant (*R*_m_*C*_m_) of 38 ms (Schmidt-Hieber et al., 2007).

The GC model is comprised of nine different regenerative and restorative conductances: fast sodium (NaF), hyperpolarization-activated cyclic-nucleotide-gated (HCN), L-type calcium (CaL), N-type calcium (CaN), T-type calcium (CaT), delayed rectifier potassium (KDR), A-type potassium (KA), big conductance (BK), and small conductance (SK) calcium activated potassium. Hodgkin–Huxley (HH) or Goldman-Hodgkin-Katz (GHK) formulations (Goldman, 1943; Hodgkin and Katz, 1949; Hodgkin and Huxley, 1952) were employed to model these voltage-and/or calcium-gated ion channels (Mishra and Narayanan, 2019). The GHK formulation was used to model calcium conductances, with intracellular and extracellular calcium concentration set at 50 nM and 2 mM, respectively. The reversal potential values for Na, K, and HCN channels were set as +55 mV, −90 mV and −30 mV, respectively. Cytosolic calcium concentration and its evolution with time was dependent on calcium current and its decay, and the mechanism was adopted from the formulation (Destexhe et al., 1993; Poirazi et al., 2003; Carnevale and Hines, 2006; Narayanan and Johnston, 2010):

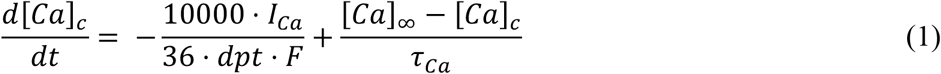

where *F* is Faraday’s constant, the calcium decay constant in GCs was given by *τ*_*Ca*_ with a default value of 160 ms, *dpt* represented the depth of the shell into which calcium influx occurred and was taken as 0.1 μm, and [*Ca*]_∞_ = 50 nM was considered as the steady state value of [*Ca*]_*c*_.

We generated 20,000 models of GC through a stochastic search from a parametric space comprised of 40 different parameters (Table 1): 38 parameters associated with nine active conductances along with 2 passive neuronal parameters. The GC models were declared valid once they fall within the range of nine physiologically constrained measurements (Table 2): input resistance (*R*_in_), sag ratio, firing rate at 50 pA (*f*_50_) and 150 pA (*f*_150_) current injection, spike frequency adaptation, action potential (AP) amplitude, AP threshold, AP half width, and fast afterhyperpolarization. The validation process resulted in 126 valid models that manifested characteristic electrophysiological properties of granule cells, but exhibited pronounced heterogeneities in channel composition and other biophysical parameters (Mishra and Narayanan, 2019). This constitutes an instance of ion-channel degeneracy (Mishra and Narayanan, 2019; Rathour and Narayanan, 2019; Goaillard and Marder, 2021; Mishra and Narayanan, 2021a) in the emergence of cellular-scale properties, and provided 126 GC models that were endowed with signature heterogeneities in their intrinsic properties. In our analyses, this population of 126 GC models from (Mishra and Narayanan, 2019) was employed as the substrate for assessing the impact of *intrinsic heterogeneities* on synaptic plasticity profiles.

**Table 1.**
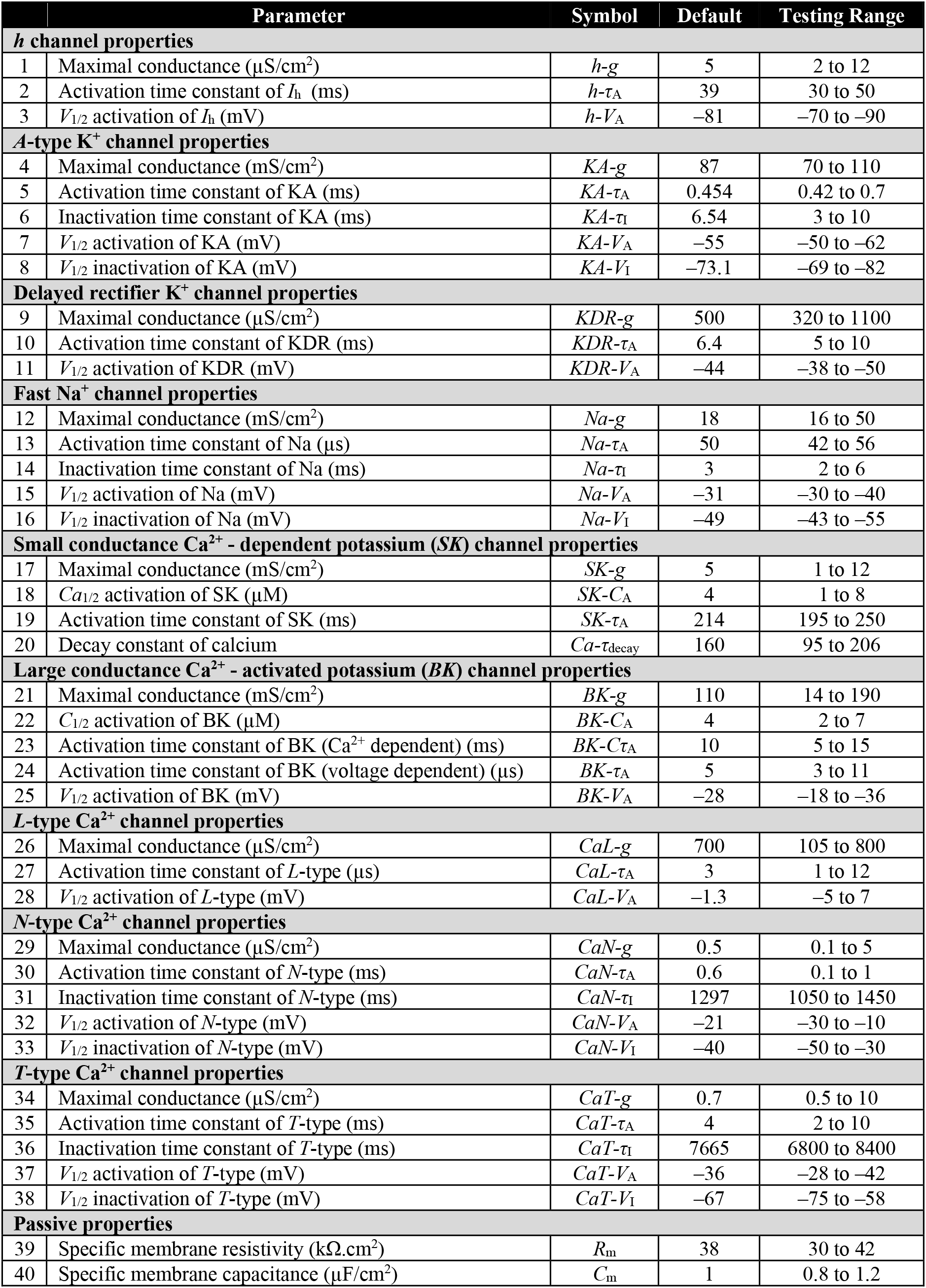
The multiple parameters and their ranges for the stochastic search employed for finding the 126 valid granule cells (Mishra and Narayanan, 2019).

**Table 2.**
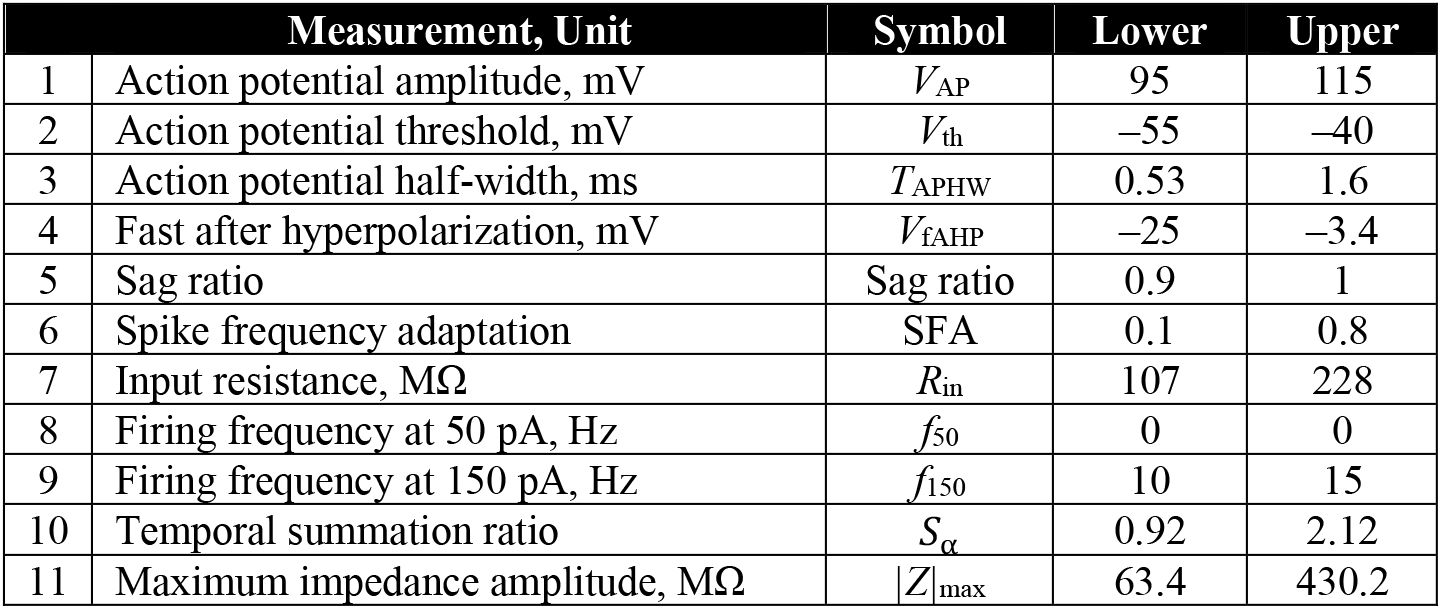
Electrophysiological bounds for the multiple objectives, defining characteristic granule cell measurements, of the stochastic search procedure (Mishra and Narayanan, 2019). The first 9 measurements were employed to validate the 126 intrinsically heterogeneous model neurons (Mishra and Narayanan, 2019), whereas the last two measurements were validated for the 126 models (Fig. 1) with electrophysiological bounds derived from (Mishra and Narayanan, 2020a).

### Properties and associated heterogeneities in synapses impinging on granule cell models

We modeled a canonical synapse impinging on the postsynaptic GC neuron as two co-localized excitatory synaptic receptors: α-amino-3-hydroxy-5-methyl-4-isoxazolepropionic acid (AMPA) receptor (AMPAR) and N-methyl-D-aspartate (NMDA) receptor (NMDAR) with an NMDA: AMPA ratio value of 1.5. The current through AMPAR and NMDAR as a function of voltage and time are modeled using the GHK formulation (Goldman, 1943; Hodgkin and Katz, 1949) as a sum of current generated by sodium and potassium ions (Narayanan and Johnston, 2010; Honnuraiah and Narayanan, 2013; Anirudhan and Narayanan, 2015):

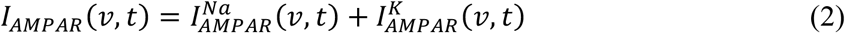

where,

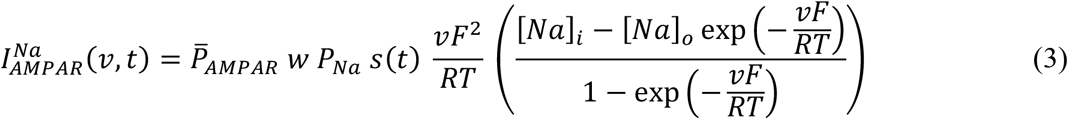

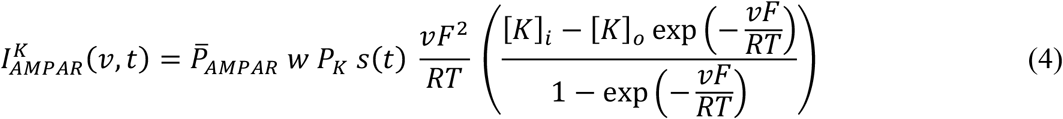

Here, 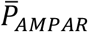 represents the maximum permeability of the receptor, also used as a synaptic parameter to incorporate synaptic heterogeneity. *w* represents the synaptic weight parameter that would be updated and monitored as a function of time to quantify positive and negative weight changes based on the plasticity protocol (see below). The default value of initial weight, *w*_*init*_ was set to 0.25. The sodium (*P*_*Na*_) and potassium (*P*_*K*_) permeability values were set to be equal (*P*_*Na*_: *P*_*K*_=1:1) based on experimental observations. The default values for intracellular and extracellular concentration (mM) of specific ions were: [*Na*]_*i*_ = 18, [*Na*]_*o*_ = 140, [*K*]_*i*_ = 140, [*K*]_*o*_ = 5 which led to equilibrium potential of +55 mV and −90 mV for Na and K, respectively. *s*(*t*) guides the kinetics of AMPA current as represented using the two-exponential formulation:

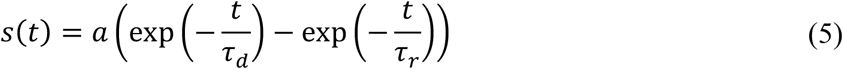

where *a* represents normalization constant so that 0 ≤ *s*(*t*) ≤ 1. *τ*_*r*_ and *τ*_*d*_ denote the rise and decay time constants associated with AMPA receptor with values of 2 ms and 10 ms, respectively. *Synaptic heterogeneities* were introduced into the population of models by altering the permeability value of 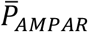.

The current through NMDA receptor depended on sodium, potassium and calcium ions and was modeled as follows using the GHK formulation:

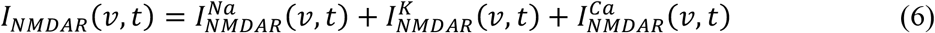

where,

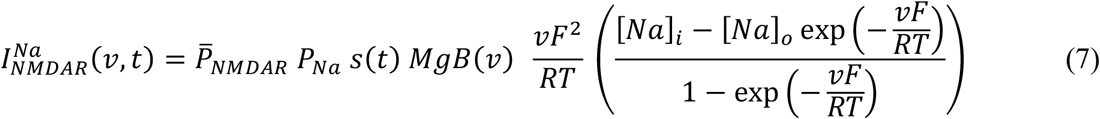

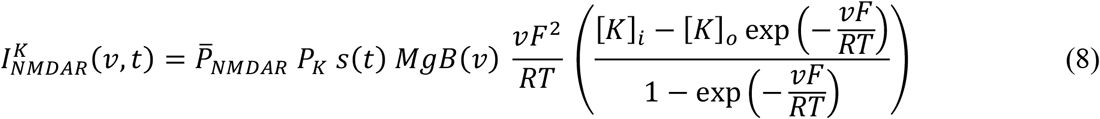

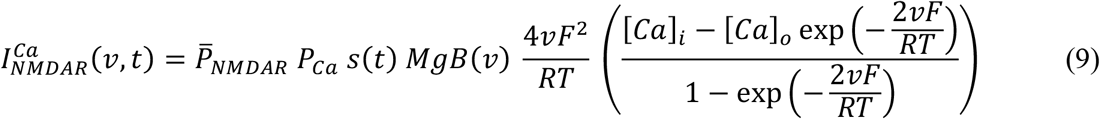

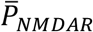 denotes the maximum permeability of the NMDA receptor and was defined as the product of 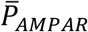, *w*_*init*_ and the value of NMDA:AMPA ratio. The permeability ratios of three ions for NMDAR is set as *P*_*Ca*_: *P*_*Na*_: *P*_*K*_ = 10.6:1:1 (Mayer and Westbrook, 1987; Canavier, 1999). The *s*(*t*) function was same as for AMPAR with *τ*_*r*_ = 5 ms and *τ*_*d*_ = 50 ms. The concentration values in mM are: [*Na*]_*i*_ = 18, [*Na*]_*o*_ = 140, [*K*]_*i*_ = 140, [*K*]_*o*_ = 5 [*Ca*]_*i*_ = 100 × 10^−6^, and [*Ca*]_*o*_ = 2. *MgB*(*v*) refers to the dependence of NMDAR currents on magnesium (Jahr and Stevens, 1990). The current through NMDAR did not undergo plasticity.

### Heterogeneities in structural properties of the granule cell population

*Structural heterogeneities*, mediated by the expression of adult neurogenesis in the DG, were incorporated into the GC model population by subjecting the mature set of 126 valid models to structural plasticity. Specifically, the reduction in dendritic arborization and in the overall number of channels expressed in immature neurons (Aimone et al., 2014) was approximated by a reduction in the diameter of the model neuron, using *R*_in_ as the measurement to match with experimental counterparts (Mishra and Narayanan, 2019). Electrophysiologically, *R*_in_ of mature and immature cells have been measured to be in the ~100–300 MΩ and ~3–6 GΩ ranges, respectively (Schmidt-Hieber et al., 2004; Overstreet-Wadiche et al., 2006b; Pedroni et al., 2014; Heigele et al., 2016; Mishra and Narayanan, 2020a, 2021a). Reducing the diameter of the models in neural population increased neuronal excitability, reflecting as increased *R*_in_ and increased firing rate. To assess the impact of structural heterogeneities on synaptic plasticity profiles, we varied the diameter of the 126 neurons in the model population from 1 μm to 63 μm. A diameter range of 2–9 μm yielded *R*_in_ values that matched the experimental 3–6 GΩ range for immature neurons and was considered representative of the immature neuronal models (Mishra and Narayanan, 2019).

### Intrinsic measurements

The 126 GC models were selected based on the nine physiological measurements employed to characterize the valid GC population (Table 2; (Mishra and Narayanan, 2019)). In addition to these, we introduced two more sub-threshold measurements (impedance amplitude and temporal summation ratio) to test the robustness of these intrinsically heterogeneous models (Fig. 1*C–D*), and to compare their role in regulating plasticity profiles. Specifically, we employed input resistance (*R*_in_), firing frequency to pulse current injections, sag, impedance amplitude and temporal summation as intrinsic measurements in relating them to plasticity profiles. *R*_in_ was measured as the slope of a linear fit to the *I-V* plot. The *I-V* plot was obtained by plotting the steady state value of voltage response as a function of 11 different current pulses where the amplitude varied from −50 pA to +50 pA in steps of 10 pA (Fig. 1*B*). As GC models with lower diameters manifested high excitability, *R*_in_ was computed in response to hyperpolarizing current pulses ranging from −50 pA to 0 pA in steps of 10 pA, to avoid spike generation. To characterize the impedance amplitude profiles of these models, we injected chirp stimulus, a frequency-dependent current input with linearly increasing frequency from 0–15 Hz in 15 s of constant amplitude (Mishra and Narayanan, 2020a). The impedance profile *Z*(*f*) was computed as the ratio of the Fourier transform of voltage response to the Fourier transform of chirp current as a function of frequency (Fig. 1*C*). The impedance amplitude profile was calculated as:

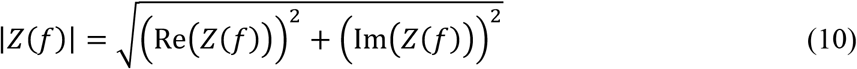

where Re(*Z*(*f*)) and (*Z*(*f*)) refer to the real and imaginary parts of the impedance *Z*(*f*), respectively, as functions of frequency *f*. The maximum value of impedance across all frequencies was measured as the maximum impedance amplitude (|*Z*|_max_).

**Figure 1:**
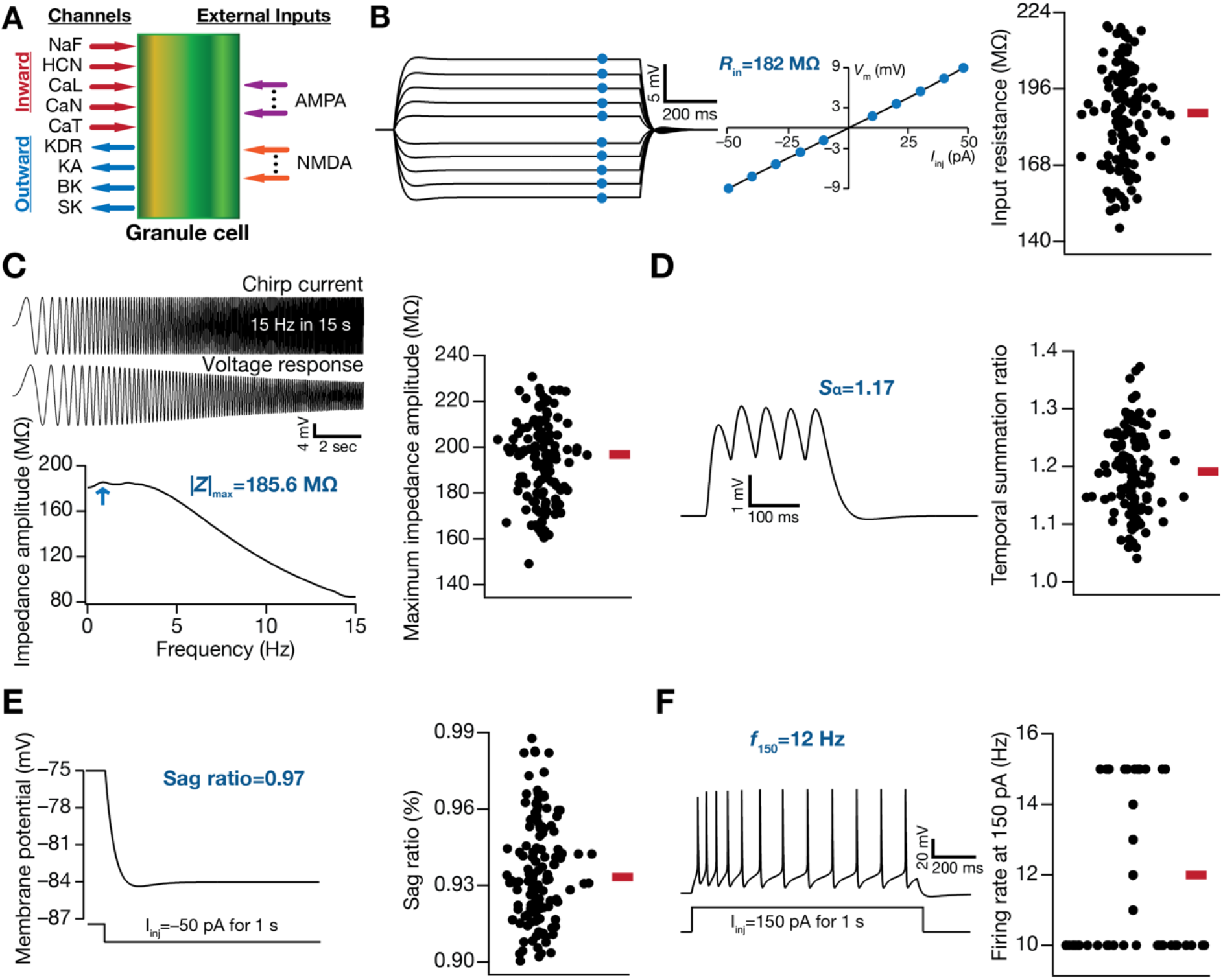
Model components of dentate gyrus granule cells and illustration of intrinsic heterogeneities across different physiological measurements. (A) Conductance-based single compartmental model of granule cell expressing different inward and outward voltage-dependent ion-channel currents, receiving excitatory inputs modeled as AMPA and NMDA receptor currents. (B–F) Different intrinsic physiological measurements employed to define the valid population of GC models (*N*_GC_ = 126). (B) *Left*, voltage traces in response to current pulses of amplitude −50 pA to +50 pA, in steps of 10 pA. *Right*, input resistance (*R*_in_), calculated as the slope of the *V–I* curve obtained by plotting the steady-state voltage responses against injected current amplitudes. (C) *Top*, A chirp current stimulus of 50 pA peak-to-peak amplitude with linearly increasing frequency from 0 to 15 Hz in 15 s depicted along with the respective voltage response. *Bottom*, the impedance amplitude profile obtained from the chirp current and voltage response shown above. (D) *Left*, Voltage response of a GC model to current input comprised of 5 α-EPSCs arriving at 20 Hz, to compute temporal summation ratio (*S*_*α*_). *S*_*α*_ is the ratio of the voltage amplitude in response to the 5^th^ α-EPSC to that of the 1^st^ α-EPSC. (E) *Left*, membrane potential in response to 50 pA hyperpolarizing current pulse to calculate sag ratio. Sag ratio is the ratio between the steady-state voltage response and the peak voltage response. (F) *Left*, firing pattern and firing rate in response to the 150 pA depolarizing current pulse of 1 s duration. Across all panels in B–F, the *right* panels show beeswarm plots depicting heterogeneities in the respective measurement across all 126 models. The heterogeneous population of 126 GC models employed here are from (Mishra and Narayanan, 2019), with additional characterization involving new intrinsic measurements added to the validation process.

Temporal summation ratio (*S*_*α*_) was computed by injecting current pattern following the *I*_*α*_(*t*) = *I*_max_*t* exp(−*αt*) formulation, where α = 0.1 ms^−1^. Five such current pulses were injected into the neuron with 50 ms interval between them, together resulting in a response consisting of five α excitatory postsynaptic potentials (α–EPSPs). The ratio of amplitude of last to first EPSP (E_last_/E_first_) was defined as the temporal summation ratio, *S*_*α*_ (Fig. 1*D*). Sag ratio was computed as the ratio between the steady-state voltage deflection to the peak voltage deflection from *V*_RMP_ in response to hyperpolarizing current pulse of 50 pA injected for a period of 1 s. The firing property of GC models was characterized by computing the firing rate in response to a current pulse of 100 pA (*f*_100_) or 150 pA (*f*_150_) for 1 s (Fig. 1*F*).

### Synaptic plasticity protocols and weight evolution

The synaptic weight parameter *w* governing current through AMPAR depended on the intracellular calcium concentration as follows, based on the calcium control hypothesis (Shouval et al., 2002):

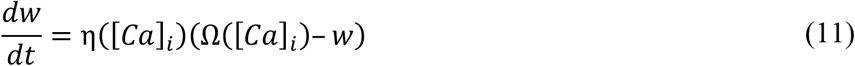

where η([*Ca*]_*i*_) represents learning rate dependent on calcium concentration, which is inversely related to learning time constant *τ*([*Ca*]_*i*_) as follows:

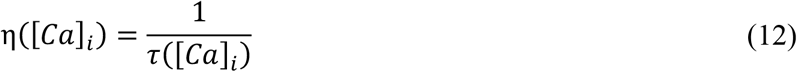

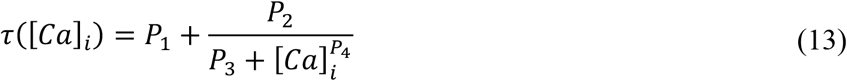

where *P*_1_ = 1 s, *P*_2_ = 0.1 s, *P*_3_ = *P*_2_ × 10^−4^, *P*_4_ = 3. These values when substituted in the equation (12) sets the learning time constant to ~ 3 hrs when [*Ca*]_*i*_ is ~ 0. Ω([*Ca*]_*i*_), the function that governed the calcium-dependent weight update mechanism, was defined as (Shouval et al., 2002):

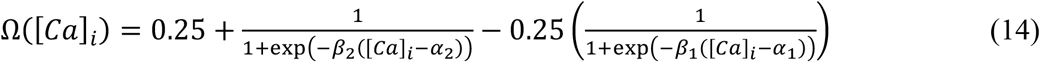

where *α*_1_ = 0.35, *α*_2_ = 0.55, *β*_1_ = *β*_2_ = 80. For all the weight update equations [*Ca*]_*i*_ was set as the deflection from the resting value of [*Ca*]_*i*_.

Using this framework, we analyzed the direction and strength of plasticity in *w* using two well-established synaptic plasticity protocols in DG neurons: the 900-pulses protocol with varying induction frequencies (Wang et al., 1997; Koranda et al., 2008; Kobayashi et al., 2013) and the theta burst stimulation (TBS) protocol (Greenstein et al., 1988; Pavlides et al., 1988; Shors and Dryver, 1994; Beck et al., 2000; Davis et al., 2004; McHugh et al., 2007; Larson and Munkacsy, 2015). The 900-pulses protocol involved synaptic stimulation made up of 900 pulses at various induction frequencies (*f*_i_ spanning 0.5–25 Hz), an experimentally and computationally well-established BCM-like (Bienenstock et al., 1982) plasticity protocol across different neurons including DG granule cells (Dudek and Bear, 1992; Wang et al., 1997; Shouval et al., 2002; Johnston et al., 2003; Koranda et al., 2008; Narayanan and Johnston, 2010; Cooper and Bear, 2012; Ashhad and Narayanan, 2013; Honnuraiah and Narayanan, 2013; Kobayashi et al., 2013; Anirudhan and Narayanan, 2015). The evolution of synaptic weight (equation 11) was monitored and the final weight at the end of the induction protocol was plotted as a function of the induction frequency (Fig. 2*B*). The percentage difference between this final weight and the initial weight (0.25) was plotted against the induction frequency of the stimulus pulses to obtain the synaptic plasticity profile (Fig. 2*C*) as a function of induction frequency (Shouval et al., 2002; Narayanan and Johnston, 2010; Honnuraiah and Narayanan, 2013; Anirudhan and Narayanan, 2015). The induction frequency at which this plasticity profile transitioned from depression to potentiation was defined as the modification threshold (Fig. 2*C*), θ_*m*_ (Dudek and Bear, 1992; Shouval et al., 2002; Johnston et al., 2003; Narayanan and Johnston, 2010; Cooper and Bear, 2012; Ashhad and Narayanan, 2013; Honnuraiah and Narayanan, 2013). We also employed percentage changes in *w* with *f*_*i*_=1 Hz (Δ*w*_1_) and *f*_*i*_=10 Hz (Δ*w*_10_) for quantifying synaptic plasticity profiles (Fig. 2*C*). The computational complexity of this process was enormous, especially in the face of three different forms of heterogeneities, and given that the construction of each profile required stimulating the synapses with 900 pulses for each of the 50 induction frequencies (*f*_*i*_ spanning 0.5–25 Hz; 0.5 Hz increment) for each of the 126 models, across several synaptic permeability and diameter values.

**Figure 2:**
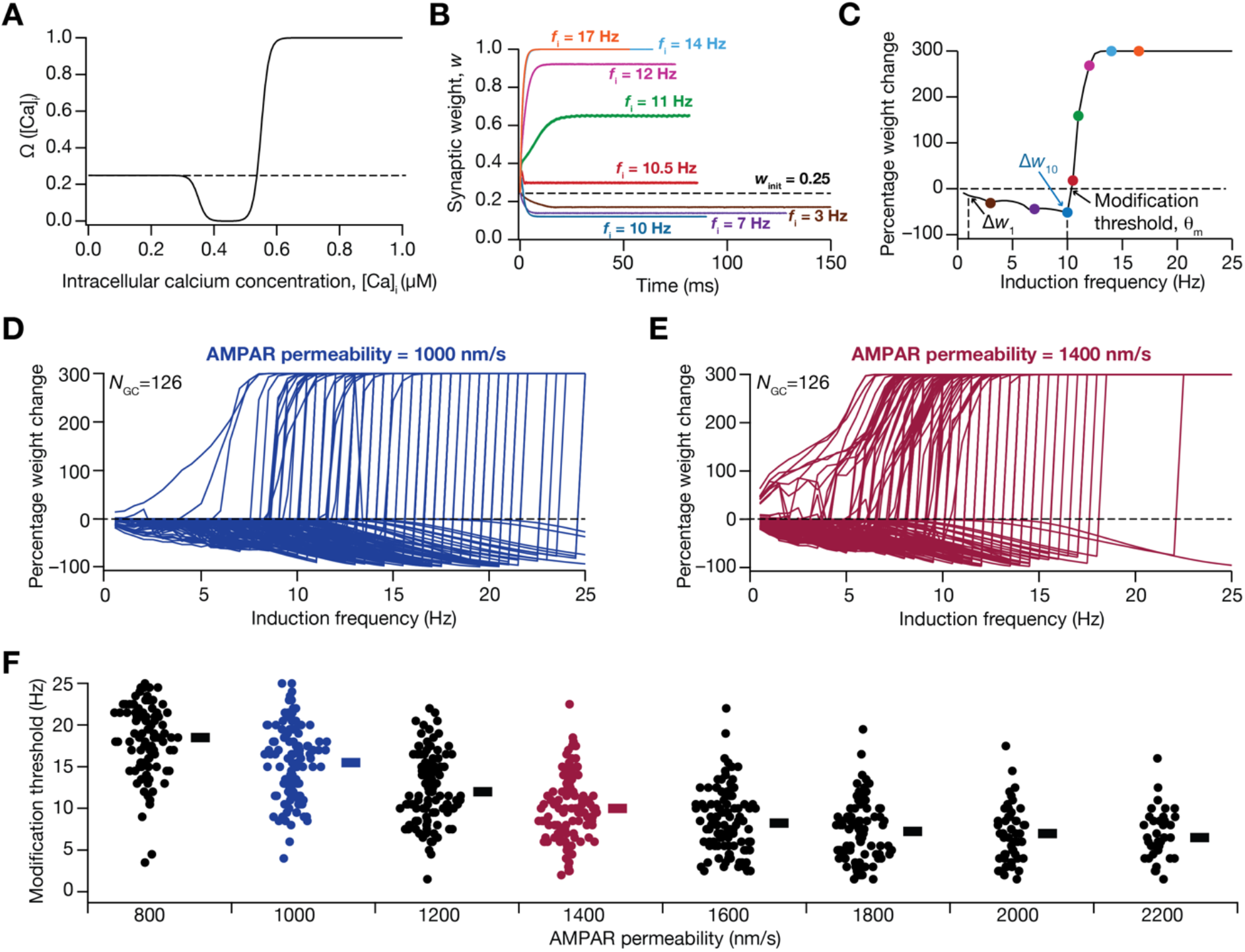
Intrinsic heterogeneities in the granule cell population translates to heterogeneities in their BCM-like synaptic plasticity profiles, when synaptic properties were fixed across models. (A) Plot of the Ω-function based on the calcium control hypothesis that regulates level of plasticity as a function of intracellular Ca^2+^ concentration (Eq. 11). (B) Evolution of synaptic weight as a function of time, obtained by employing 900-pulses protocol of different induction frequencies in a granule cell model. Note that all plots initialize at *w*_*init*_ = 0.25 and evolve to reach their respective steady state value. The duration of each plot spans 900 pulses at the specified induction frequency *f*_i_. (C) BCM-like synaptic plasticity profile obtained by plotting the percentage change in synaptic weight parameter after stimulation with 900-pulses of different induction frequencies ranging from 0.5 to 25 Hz. The color-coded points correspond to the different induction frequencies shown in panel B. Arrows point to θ_m_, Δ*w*_1_ and Δ*w*_10_. Δ*w*_1_ and Δ*w*_10_ represent the change in synaptic weight value for induction frequencies of 1 Hz and 10 Hz, respectively; θ_m_, the modification threshold, is the induction frequency at which the plasticity profile switches from inducing LTD to inducing LTP. (D–E) Same as C, for all the 126 GC models for two different values of AMPAR permeability: 1000 nm/s (D) and 1400 nm/s (E). (F) Beeswarm plots of modification threshold for all GC models, for different values of AMPAR permeability. Note that with specific values of AMPAR permeability, there were models that did not manifest a θ_m_ in the tested range of frequencies, thus resulting in lesser number of models for each AMPAR permeability values (*N*=100, 121, 121, 120, 112, 94, 63, 39 left to right).

For TBS, the synapse was stimulated with a burst of 5 action potentials at 100 Hz, and this burst was repeated 150 times at 200 ms interval (theta frequency) each (Fig. 5*A*). This was done to achieve steady state values for [*Ca*]_*i*_ and *w* (Ashhad and Narayanan, 2013). The percentage change in *w* at the end of this protocol in comparison to its initial value (*w*_init_ = 0.25) was employed to quantify plasticity induced with TBS. For both plasticity induction protocols, we employed a spike train generator as an input source to mimic pre-synaptic activity.

### Computer simulations and analysis

We employed NEURON as the simulation environment (Carnevale and Hines, 2006) for executing all the simulation at *V*_RMP_ (−75 mV) with fixed temperature set at 34°C. We used the integration time step of 25 μs except for simulations involving 900-pulses protocol, where a variable time step method was employed to efficiently solve the associated differential equations with lower computational time. All data analyses were performed using custom-built software under the Igor-Pro programming environment (Wavemetrics Inc., USA). To avoid ambiguities arising from reporting merely the summary statistics (Marder and Taylor, 2011; Rathour and Narayanan, 2019), we have reported all the data points with their respective ranges to represent the heterogeneities associated with our analysis and results. As we have employed Pearson correlation coefficient for pairwise scatter plots, qualitative descriptions on the strength of correlation coefficient values (weak *vs.* strong) were adopted from the definitions provided in (Evans, 1996).

## RESULTS

We employed a physiologically realistic conductance-based population of GC models (*N*_GC_=126), endowed with *intrinsic heterogeneities* and expressing ion-channel degeneracy at the cellular-scale (Mishra and Narayanan, 2019), to assess the impact of neural heterogeneities on synaptic plasticity profiles. In this population, we introduced *synaptic heterogeneities* by altering afferent synaptic strength, and *structural heterogeneities* by changing the surface area of the model population. We employed two well-established synaptic plasticity protocols, namely the BCM-like 900-pulses protocol with different induction frequencies and the theta burst stimulation (TBS) protocol, to examine the impact of these three forms of neural heterogeneities in the regulation of synaptic plasticity rules in DG granule cells. We present results obtained through systematic incorporation of these forms of heterogeneities, both independently and synergistically, into a physiologically validated GC model population.

### GC models showed robustness for non-validated measurements and manifested heterogeneities in intrinsic measurements

The 126 GC models employed in this study were previously validated based on nine different characteristic electrophysiological signatures of DG granule cells (Mishra and Narayanan, 2019). Prominent among these measurements are input resistance (*R*_in_, range 140–225 MΩ, Fig. 1*B*), sag ratio (range 0.9–1, Fig. 1*E*) and firing rate at 150 pA (range 10–15 Hz, Fig. 1*F*), which manifested heterogeneities. In addition to these, here we characterized two more experimentally obtained sub-threshold measurements of excitability to assess their relationship to the induction of synaptic plasticity: impedance amplitude and temporal summation ratio (Fig. 1*C–D*). Whereas temporal summation of postsynaptic potentials constitutes an important measurement that governs calcium influx, and thereby synaptic plasticity (Nolan et al., 2004; Narayanan and Johnston, 2010), impedance is a measure of excitability for time-varying signals (Narayanan and Johnston, 2008). Although the 126 GC models were not validated against these two measurements, we found that these measurements in the models were within the range of their electrophysiological counterparts (Mishra and Narayanan, 2020a). Specifically, maximum impedance amplitude (|*Z*|_max_) in the model population ranged from 149.1 MΩ to 230.8 MΩ (mean ± SEM: 194.2 ± 1.6; *N*_GC_=126; Fig. 1*C*), which was within the measured electrophysiological range (Mishra and Narayanan, 2020a) of 63.4–430.2 MΩ (mean ± SEM: 176.9 ± 5.3; *N*=172). The temporal summation ratio in the model population ranged from 1.04–1.37 (mean ± SEM: 1.19 ± 0.006; *N*_GC_=126; Fig. 1*D*), which was within the measured electrophysiological range (Mishra and Narayanan, 2020a) of 0.92–2.12 (mean ± SEM: 1.33 ± 0.015; *N*=133). Apart from providing two additional intrinsic measurements, these analyses showed that these intrinsic properties manifested heterogeneities in the model population (Figs. 1*C–D*) and provided further validation of the population of models in terms of their ability to match with characteristic signatures of GCs. In addition, the parameters (spanning active and passive neural properties; Table 1) underlying these 126 models manifested considerable heterogeneities, thus providing a layer of biophysical heterogeneities in the GC population.

### Intrinsic heterogeneity resulted in heterogeneities in BCM-like plasticity profiles when models received identical synaptic inputs

To understand the impact of intrinsic heterogeneities on emergence of plasticity profiles, we first employed the well-established BCM-like 900 pulses protocol and constructed the synaptic plasticity profile, spanning different induction frequencies (*f*_*i*_), of each GC model. The stimuli comprised of 900 synaptic stimulations impinging on a synapse on each model neuron, at different induction frequencies ranging from 0.5 to 25 Hz. The synaptic stimulation was allowed to activate a synapse endowed with co-localized AMPAR and NMDAR, with identical values for receptors’ densities and properties across all GC models. Activation of these receptors resulted in influx of calcium into the cytosol, through NMDARs and voltage-gated calcium channels (VGCC) expressed in the models, with the strength and the dynamics of calcium evolution depending upon the induction frequency and the specific model under consideration. Although the synaptic properties and stimulation protocols were identical across models, the cytosolic calcium influx would be model-dependent because of the differential expression of VGCCs across models. The influx of calcium, in turn, affected the weight parameter (*w*) associated with the AMPARs, following the calcium control hypothesis (Fig. 2*A*; Equation 11). We monitored the temporal evolution of the weight parameter and recorded the final value at the end of protocol (after 900 pulses of the specific *f*_*i*_) for each induction frequency (Fig. 2*B*). To obtain the synaptic plasticity profile, the percentage weight change in *w* was computed from its final value for each *f*_*i*_ with respect to the initial value (*w*_init_=0.25) and was plotted as a function of *f*_*i*_ (Fig. 2*C*). Consistent with experimental results from DG granule cells (Wang et al., 1997; Koranda et al., 2008; Kobayashi et al., 2013) and with the Ω-function that governs synaptic plasticity (Fig. 2*A*), we found that lower and higher values of *f*_*i*_ yielded depression and potentiation, of AMPAR weight, respectively (Fig. 2*C*). The induction frequency at which the synaptic plasticity profile transitioned from depression to potentiation was termed the modification threshold θ_O_ (Bienenstock et al., 1982; Shouval et al., 2002; Narayanan and Johnston, 2010; Honnuraiah and Narayanan, 2013; Anirudhan and Narayanan, 2015).

To understand how and to what extent intrinsic heterogeneities impact the evolution of synaptic profiles and modification threshold, we obtained plasticity profiles for all the 126 GC models with *identical* structural and synaptic properties. We found that heterogeneities in intrinsic properties of these models resulted in heterogeneities in the BCM-like plasticity profiles (Fig. 2*D–E*) as well as in the associated modification thresholds (Fig. 2*F*). We repeated these analyses for different values of baseline synaptic strength (defined as receptor permeability, 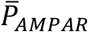, within the GHK formulation for AMPARs) to explore the association of the fixed synaptic parameter to intrinsic heterogeneities in altering the plasticity profiles (Fig. 2*D–F*). We observed a graded reduction in the modification threshold (Fig. 2*F*), implying a leftward shift in the BCM-like plasticity profile, with increase in the baseline synaptic strength. This is to be expected because with increased synaptic strength, the postsynaptic depolarization and consequently the cytosolic calcium influx are higher, thus allowing the plasticity profile to transition to synaptic potentiation at lower induction frequencies (Shouval et al., 2002; Narayanan and Johnston, 2010). As a consequence of intrinsic heterogeneities across models and such leftward shifts in plasticity profile, there was also an increase in the number of models that manifested no synaptic depression (within the tested range of *f*_*i*_) with increases in baseline synaptic strength. Together, these results demonstrated that the expression of intrinsic heterogeneities led to heterogeneities in synaptic plasticity profiles, when structural and synaptic properties across models were identical.

### Weak pair-wise correlations between intrinsic and plasticity-profile measurements

How are the different intrinsic measurements defining the 126 GC models related to the measurements employed to quantify the synaptic plasticity profiles? Does a specific range of physiological sub- or suprathreshold properties determine the synaptic plasticity measurements or are they independent of each other? Does the synaptic permeability parameter play any role in defining the relationship between these intrinsic and synaptic plasticity measurements? To address these questions, we plotted intrinsic measurements against measurements related to the synaptic plasticity profiles for all the 126 GC models, for two different values of baseline synaptic strength (Fig. 3). We employed five intrinsic measurements (Fig. 1), namely input resistance (*R*_in_), temporal summation (*S*_*α*_), sag ratio, maximum impedance amplitude (|*Z*|_max_), and firing rate at 150 pA (*f*_150_). Three measurements related to the synaptic plasticity profile were employed, namely percentage weight changes at *f*_*i*_=1 Hz (Δ*w*_1_) and 10 Hz (Δ*w*_10_), modification threshold (θ_*m*_). We first plotted the pair-wise scatter plots between the intrinsic and the synaptic plasticity measurements spanning all 126 GC models and calculated the Pearson’s correlation coefficient for these pairwise scatter plots (Fig. 3). We found weak pairwise correlation coefficients (−0.4 < *R* < 0.4) across all the pairs, when synaptic plasticity measurements were computed with two different values of baseline synaptic strength (Fig. 3*A–B*). These results suggest that intrinsic excitability and temporal summation are not sufficiently strong to impose specific plasticity profiles on model synapses across the heterogeneous population of models, and that several other mechanisms govern the emergence of these plasticity profiles.

**Figure 3:**
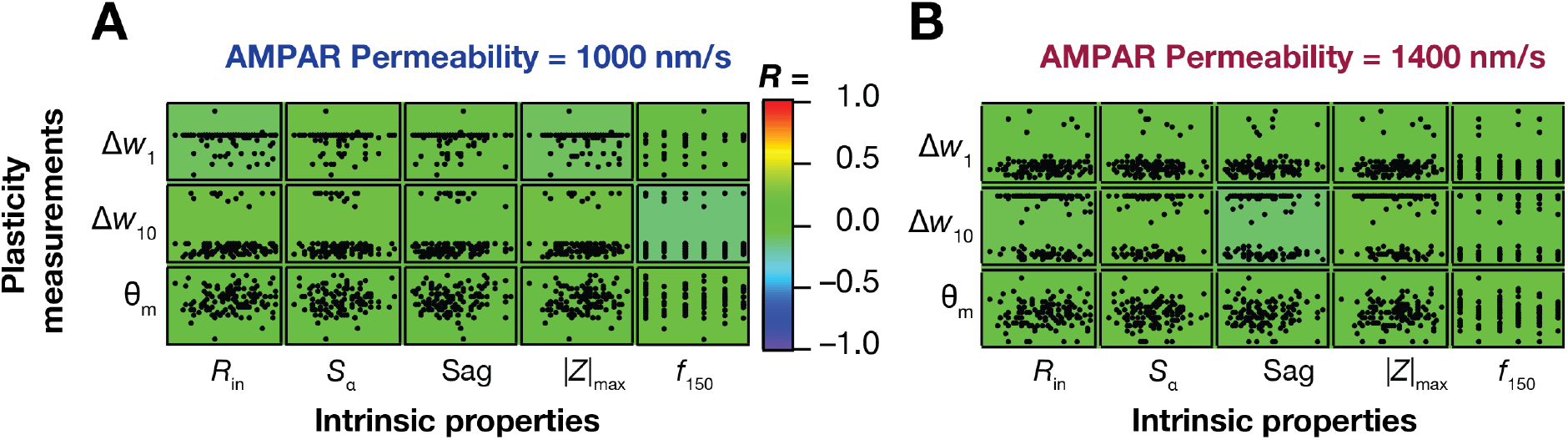
Weak pair-wise correlations between measurements of synaptic plasticity and intrinsic properties in the heterogenous GC model population. (A–B) Pair-wise scatter plot matrices between three plasticity measurements: percentage weight change at 1 Hz (Δ*w*_1_), 10 Hz (Δ*w*_10_) and the modification threshold (θ_m_) along the vertical axes, and five intrinsic measurements: *R*_in_, *S*_*α*_, *f*_150_, |*Z*|_max_, and Sag on the horizontal axes. Synaptic plasticity measurements were obtained for baseline AMPAR permeability values of 1000 nm/s (A) and 1400 nm/s (B). The scatter plot matrices are overlaid on the respective color-coded values of correlation coefficients (*R*).

### Synergistic interactions between neuronal intrinsic properties and synaptic strength result in the emergence of similar synaptic plasticity profiles

The analyses thus far assumed the baseline synaptic strength (defined by receptor densities prior to plasticity induction) to be uniform across all valid GC models. Could synapses across these neuronal models manifest similar plasticity profiles despite the expression of pronounced heterogeneities in their intrinsic properties? As baseline synaptic strength is a known modulator of plasticity profiles (Shouval et al., 2002; Narayanan and Johnston, 2010; Anirudhan and Narayanan, 2015), could a model-dependent baseline synaptic strength allow for the emergence of similar plasticity profiles across all models?

To address these questions, we first executed an algorithm, independently for each of the 126 models, that identified the value of baseline synaptic strength 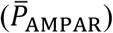 that yielded a synaptic plasticity profile with the modification threshold around 10 Hz (9.75 ≤ θ_*m*_ ≤ 10.25). Despite the considerable heterogeneities in intrinsic properties, we found that altering 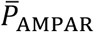 was sufficient to achieve similar synaptic plasticity profiles across all 126 models (Fig. 4*A*) with θ_*m*_ falling within the tight bound (Fig. 4*B*). The considerable heterogeneities in intrinsic properties, however, manifested in the required 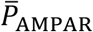 value to achieve similar plasticity profiles. The value of 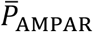 required to achieve similar plasticity profiles (referred to as threshold 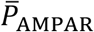) spanned a wide range (Fig. 4*C*), with the heterogeneity almost spanning an order of magnitude across models (450–3100 nm/s). Thus, although changes in 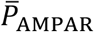 resulted in changes to the plasticity profile across models (Fig. 2*F*), *specific co-expression* of heterogeneities in synaptic (Fig. 4*C*) and intrinsic (Fig. 1) properties could result in similar plasticity profiles (Fig. 4*A–B*).

**Figure 4:**
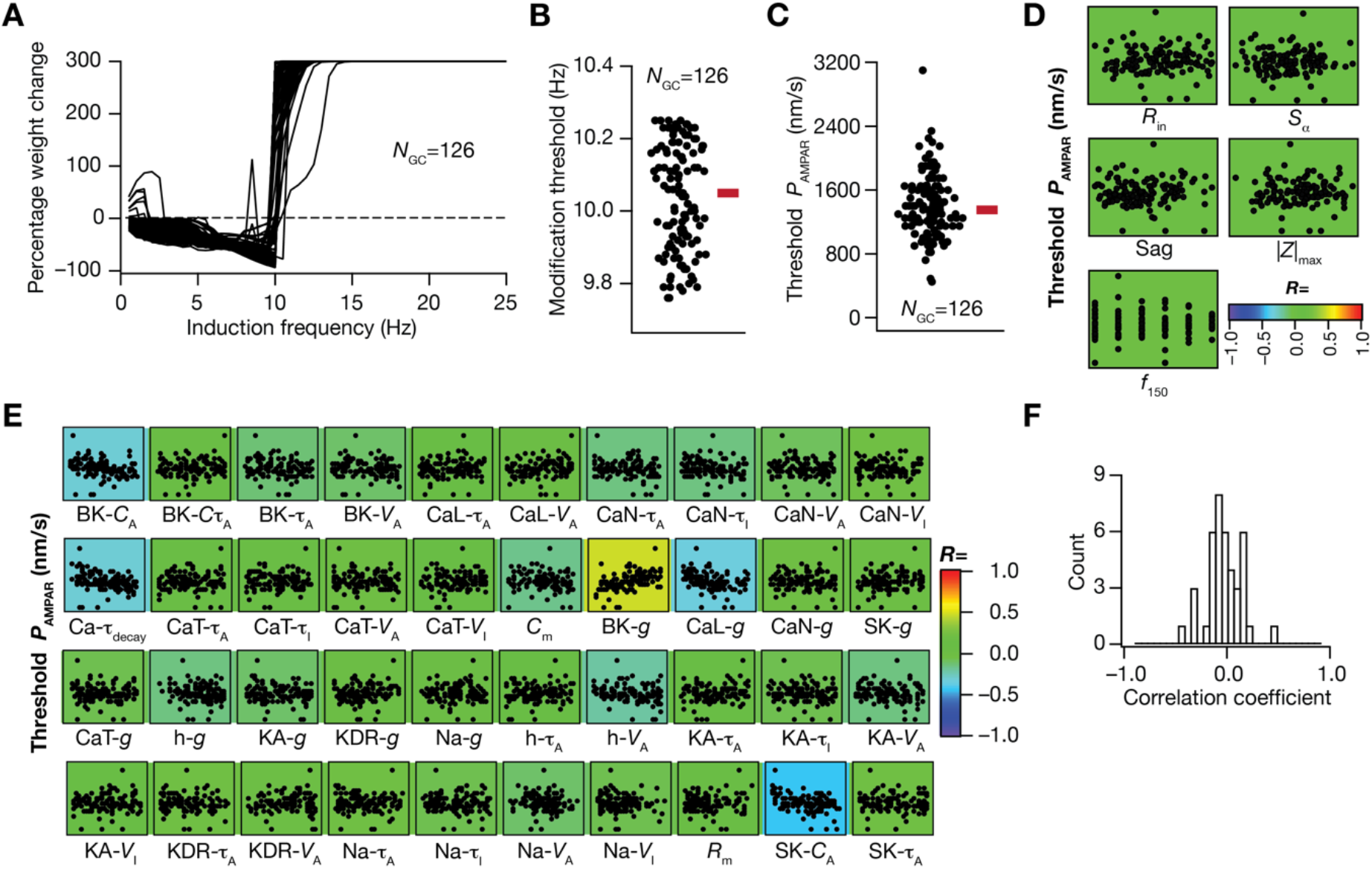
Degeneracy in the emergence of BCM-like synaptic plasticity profile resultant from synergistic interactions between heterogeneities in intrinsic and synaptic properties. (A) Similar plasticity profiles with their modification thresholds at ~10 Hz (9.75–10.25 Hz) were obtained for all 126 GC models by adjusting the synaptic strength 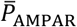 for each model. (B) Beeswarm plot shows the distribution of modification threshold around 10 Hz obtained for all the GC models. (C) Beeswarm plot representing distribution of AMPAR permeability 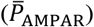 values (range: 450–3100 nm/s) required to obtain similar plasticity profile (panels A–B) across the 126 GC models. (D) Pair-wise scatter plots between AMPAR permeabilities shown in panel C and five intrinsic measurements, overlaid on respective color-coded correlation values showing weak pair-wise correlations, plotted for the 126 GC models. (E) Pair-wise scatter plots between AMPA permeability values shown in panel C and 40 different intrinsic channel parameters, overlaid on corresponding color-coded correlation values representing weak pair-wise correlations, plotted for the 126 GC models. (F) Histogram representing the distribution of correlation coefficient values depicted in panel *E*.

**Figure 5:**
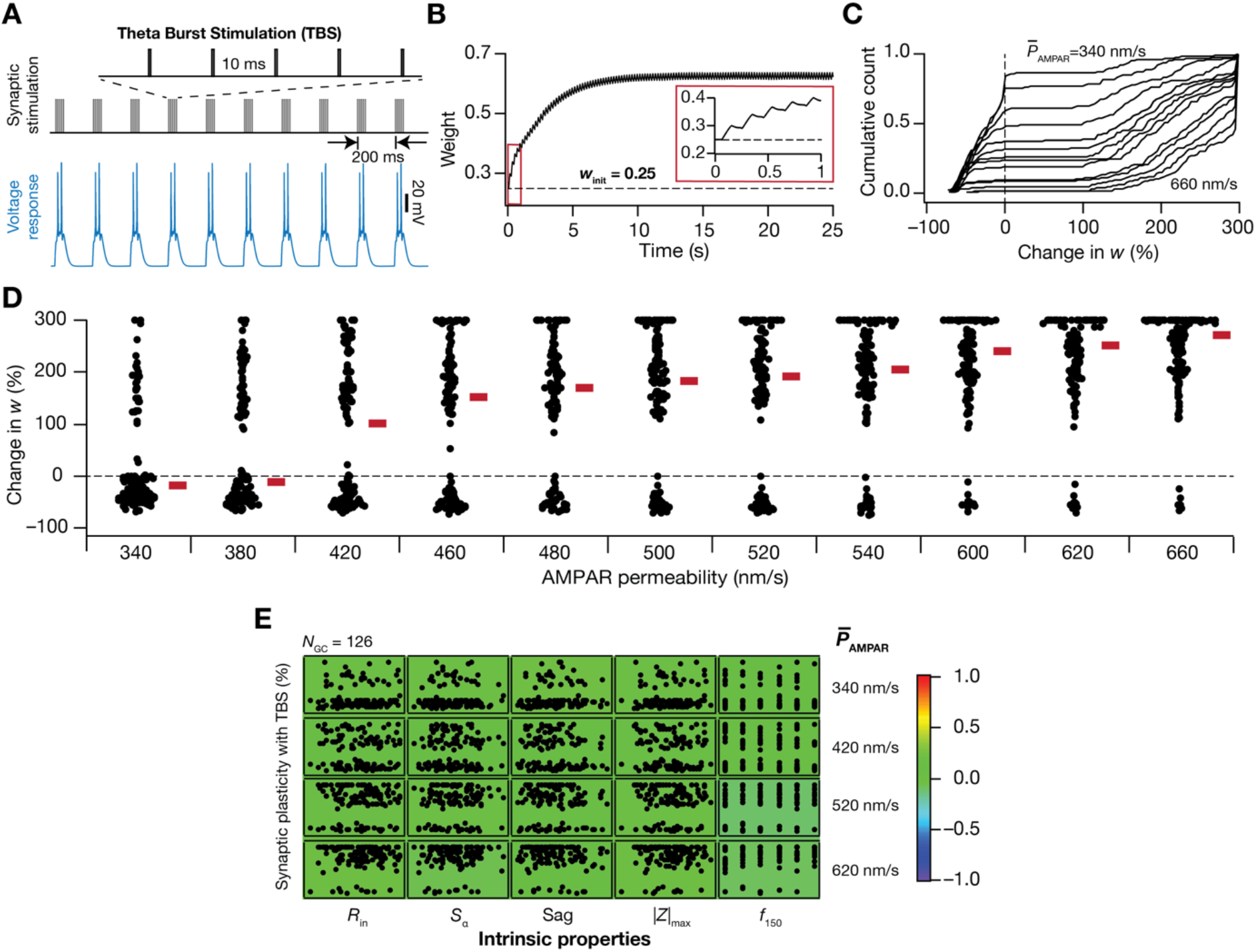
Intrinsic heterogeneities in the granule cell population translates to heterogeneities in plasticity induced by the theta burst stimulation protocol, when synaptic properties were fixed across models. (A) *Top,* Schematic of theta burst stimulation (TBS) protocol to induce synaptic plasticity. The protocol consists of bursts of stimuli with the inter burst interval set at 200 ms and each burst comprised of 5 events separated by 10 ms interval (expanded view). *Bottom*, Typical voltage response of an example GC model to TBS. (B) Plot representing evolution of AMPA receptor weight reaching an average steady-state value of 0.62 from initial weight value set to 0.25 as a function of time in response to TBS. *Inset*, Plot showing the initial portion of the weight evolution in response to five bursts of the TBS protocol. (C–D) Cumulative histogram (C) and beeswarm plots (D) showing the amount of LTP across all models with different AMPAR permeabilities, ranging from 340 nm/s to 660 nm/s. It may be noted that number of GC models undergoing LTP increases as a function of AMPAR permeability values. (E) Pair-wise scatter plots between TBS-induced change in synaptic strength and five different intrinsic properties of all GC models, plotted for different values of baseline AMPAR permeabilities. The plots are overlaid on the respective color-coded correlation coefficients values, which show weak correlations across all plots.

Did the emergence of similar plasticity profiles require strong constraints on the relationship between synaptic strength and intrinsic excitability of the models? Were there strong relationships between synaptic strength and any of the biophysical parameters that defined the models that yielded similar synaptic plasticity profiles? To address these, we first computed pairwise correlation coefficients between the 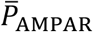 value that was required to obtain similar plasticity profiles (from Fig. 4*C*) and five intrinsic measurements of the respective models (from Fig. 1) and found them to be weakly correlated (Fig. 4*D*; −0.03 <*R*< 0.02). We next plotted pair wise scatter plot matrix between these 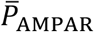 values and the 40 different intrinsic parameters that defined these 126 models to explore possible parametric dependencies. We found these pairwise correlation coefficients to be weak (−0.5 <*R*< 0.5; Fig. 4*F*), with the relatively high correlation values (|*R*|≈0.5) spanning the relationships between 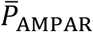 values and parameters governing cytosolic calcium dynamics (conductances of *L*-type calcium and BK channels; parameters governing calcium-dependent activation of BK and SK channels, the decay time constant of cytosolic calcium). Together, these results demonstrate that neither the intrinsic properties (spanning sub- and supra-threshold intrinsic excitability and temporal summation) nor biophysical parameters (spanning passive and active properties) are sufficient to impose strong constraints on the synaptic strength required for obtaining similar plasticity profiles. Granule cells express several mechanisms that could alter synaptically-driven cytosolic calcium elevation. This implies that heterogeneity-induced variation in any parameter is compensated by *synergistic interactions spanning* several other parameters (rather than recruiting strong pairwise compensations) towards achieving plasticity profile homeostasis. Importantly, these observations clearly demonstrate that disparate parametric combinations could yield similar plasticity profiles, pointing to the expression of degeneracy in the emergence of BCM-like synaptic plasticity profiles in DG granule cells.

### Heterogeneities and degeneracy in synaptic plasticity induced by theta burst stimulation in the heterogeneous granule cell population

The results thus far demonstrated that while heterogeneities in intrinsic and synaptic properties could independently translate to plasticity profile heterogeneity, they could synergistically interact to elicit similar plasticity profiles despite widespread heterogeneities in each underlying parameter. However, these observations were limited to the BCM-like synaptic plasticity profile. To understand the dependence and robustness of these conclusions on the type of induction protocol, we turned to a more physiologically relevant and well-established synaptic plasticity induction protocol: theta burst stimulation (TBS) (Fig. 5*A*). Synapses were provided TBS, and the consequent change in synaptic weight following the calcium-dependent dynamics (Equation 11) was computed as the difference of steady-state weight value from its initial weight (*w*_init_=0.25). The temporal evolution of synaptic weight in response to TBS in a representative model shows synaptic potentiation when steady-state weight value was achieved (Fig. 5*B*). We measured plasticity induction consequent to TBS with different values of baseline synaptic strength 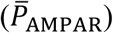 across each of the 126 intrinsically heterogeneous models.

First, for any value of 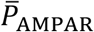, we observed pronounced heterogeneity in the magnitude and strength of TBS-induced synaptic plasticity (Fig. 5*C–D*). While synapses on certain models manifested potentiation, others showed depression for identical synapses receiving identical patterns of stimulation across models. Second, for lower values of 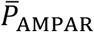 several models showed synaptic depression, whereas with higher 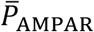, a majority manifested potentiation (Fig. 5*C–D*). Finally, there were no strong pair-wise correlations between TBS-induced synaptic plasticity and any of the several intrinsic properties of the model neurons (Fig. 5*E*). These results demonstrate that synaptic and intrinsic heterogeneities could independently alter the direction and the strength of TBS-induced synaptic plasticity, without strong pair-wise correlations between synaptic plasticity and neuronal intrinsic properties.

We next asked if we could tune each of these intrinsically heterogeneous populations of GC models to elicit similar amount of TBS-induced synaptic potentiation, despite the expression of heterogeneities. To do this, for each model, we independently executed an algorithm that searched for a 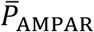 value that resulted in ~150 % TBS-induced change in synaptic strength. We found that similar amount of LTP could be obtained across all 126 intrinsically heterogeneous models (Fig. 6*A*). The 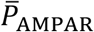 value required for yielding similar LTP, however, was heterogeneous and spanned a wide range across the 126 models (Fig. 6*B*). Thus, the specific expression of intrinsic and synaptic heterogeneities could yield similar TBS-induced LTP (Fig. 6*A–B*), despite them being independently capable of altering TBS-induced synaptic plasticity (Fig. 5). Across models, none of the five intrinsic measurements (Fig. 6*C*) or the 40 intrinsic parameters that governed the models (Fig. 6*D–E*) manifested strong pair-wise correlations with the 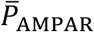 required for eliciting similar TBS-induced LTP. There were some parameters, especially those governing calcium dynamics, that manifested relatively high values of correlation coefficients with the 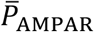 values (|*R*|≈0.5), but none of them showed strong correlations. Together, these results demonstrated the expression of degeneracy in achieving similar TBS-induced synaptic plasticity, and emphasized that intrinsic properties do not impose strong constraints on synaptic parameters towards induction of similar synaptic plasticity.

**Figure 6:**
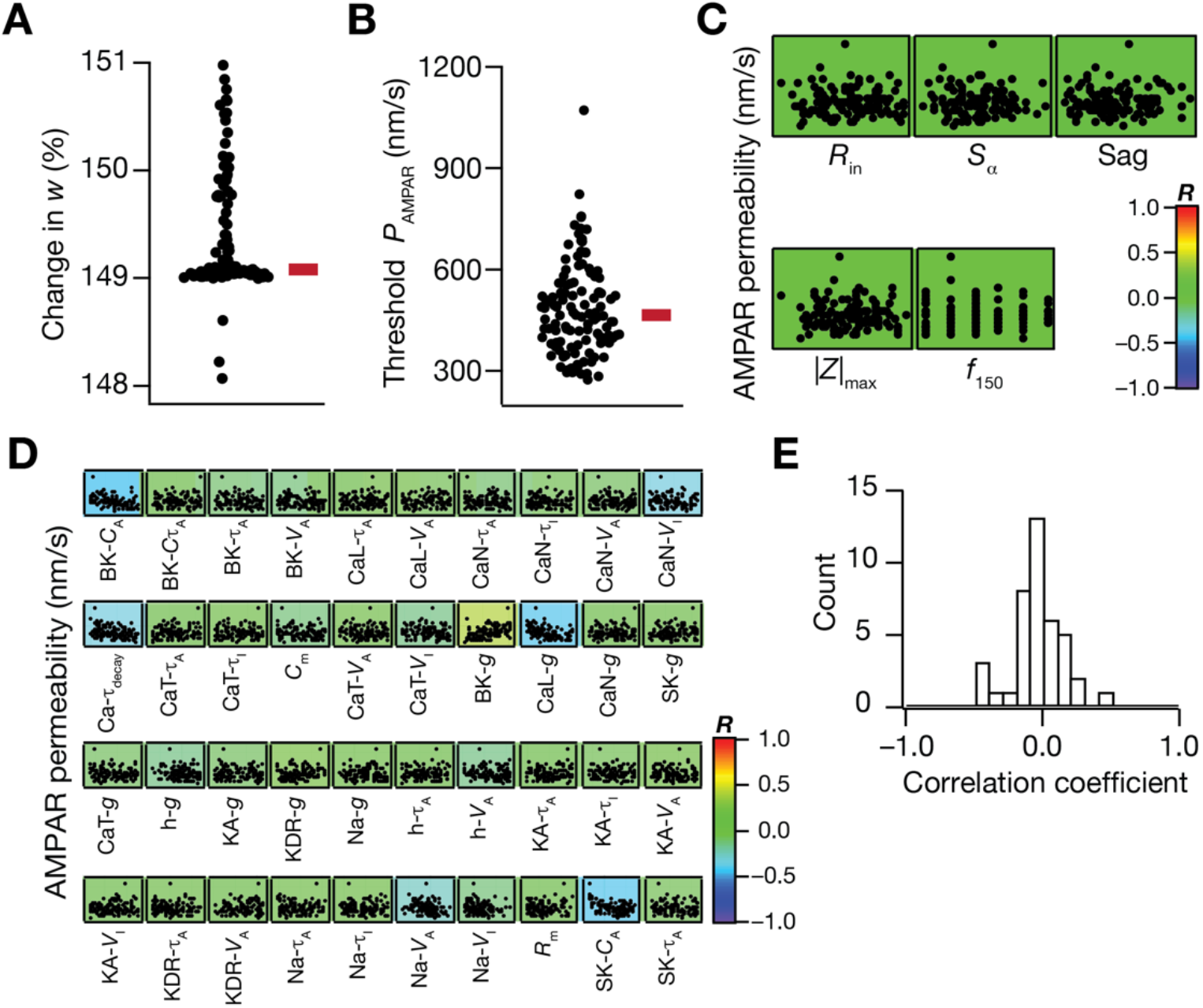
Degeneracy in eliciting the same amount of TBS-induced LTP emerges from synergistic interactions between heterogeneities in intrinsic and synaptic properties. (A) Plot showing that the same amount of LTP (~150%) is obtained in different GC models by adjusting the baseline AMPAR permeability value. (B) Beeswarm plot representing the range of AMPAR permeabilities required to obtain ~150% LTP shown in *A*. (C) Pair-wise scatter plots between permeability parameter in *B* and five intrinsic measurements of the respective models, overlaid on the respective color-coded correlation coefficients. Weak correlation values (−0.05<*R*<0.06) indicate the absence of pairwise dependency between the synaptic parameter and intrinsic measurements in the emergence of degeneracy. (D) Pair-wise scatter plots between permeability parameter in *B* and intrinsic parameters spanning all GC models. Overlaid are respective color-coded correlation values. (E) Histogram of correlation coefficients represented in *D*. Weak correlation values (−0.4<*R*<0.5) indicate lack of pairwise dependency between intrinsic and synaptic parameters in the emergence of plasticity degeneracy.

### Neurogenesis-induced age-dependent structural heterogeneity regulates the heterogeneity in plasticity profiles across intrinsically variable GC models

The dentate gyrus is endowed with adult-neurogenesis, where it takes them 4–8 weeks to fully mature and become physiologically and morphologically similar to the developmentally born neurons. Immature adult-born granule cells have reduced dendritic arborization and are highly excitable in nature with lower threshold for induction of synaptic plasticity (Schmidt-Hieber et al., 2004; Ge et al., 2007; Dieni et al., 2013; Aimone et al., 2014; Huckleberry and Shansky, 2021). To incorporate the structural heterogeneity introduced by adult neurogenesis, we independently changed the diameter (range from 1–65 μm) of 126 valid mature GCs to reflect the maturation process: 2–9 μm diameter for the immature neuronal population matching the very high input resistance found from electrophysiological studies (Schmidt-Hieber et al., 2004; Overstreet-Wadiche et al., 2006a; Overstreet-Wadiche et al., 2006b; Pedroni et al., 2014; Heigele et al., 2016; Li et al., 2017; Lodge and Bischofberger, 2019); diameters in the 60–65 μm range formed a fully mature population, based on the diameter of the base model population set at 63 μm; and intermediate diameters range 10–60 μm resulted in an age-dependent population at different maturation phases. To understand and quantify the dependence of plasticity profile on neurogenesis induced age–dependent structural heterogeneity, we first employed BCM-like 900 pulses protocol of different frequency range (0.5–25 Hz) to induce synaptic plasticity in these models. Specifically, the impact of plasticity induction was assessed in the 126 *intrinsically heterogeneous* models, with the diameter changes spanning 3–65 μm which incorporated an additional layer of *structural heterogeneity* into each of these models. A third layer of *synaptic heterogeneity* was introduced by varying the baseline AMPAR permeability 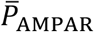 value, together providing us an experimental design that allowed us to assess the impact of all the three prominent neural-circuit heterogeneities on the synaptic plasticity profile (Fig. 7*A–E*).

**Figure 7:**
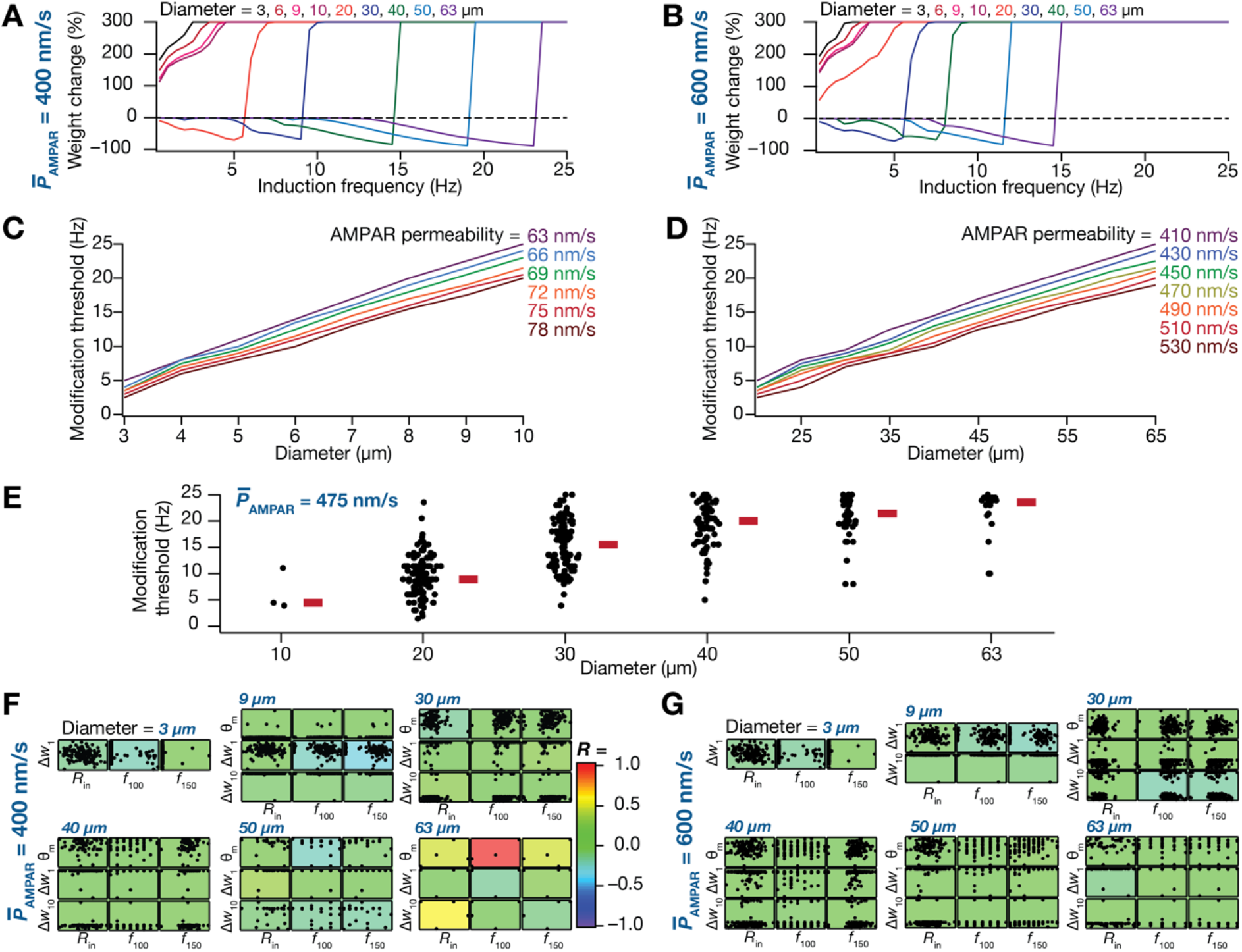
Age-dependent structural heterogeneity in GC models manifests as heterogeneity in the plasticity profiles obtained from the 900 pulses protocol. (A–B) Plasticity profiles obtained with the 900-pulses protocols with different induction frequencies, corresponding to a single GC model with different diameters for two different values of baseline AMPAR permeability: 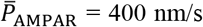 (A) and 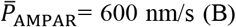 (B). (C–D) Plots of modification threshold as functions of diameter for different values of 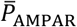 in a single GC model. Shown are plots for immature (*C*; 2–10 μm) and mature (*D*; 40–65 μm) ranges of diameters. The leftward shifts in plasticity profile observed with decreases in diameter or increases in permeability signifies lower threshold for LTP induction in the same GC model with lower diameter or higher permeability. Immature GC models undergo LTP at lower 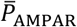 values (compare permeability ranges in panel *C vs*. panel *D*) due to their highly excitable nature. (E) Beeswarm plots showing the distribution of modification threshold as a function of diameters across all GC models for a fixed 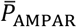 value of 475 nm/s. Note that the modification threshold did not fall within the tested range of induction frequencies for different models at different diameter values, thus resulting in different number of points for each diameter value (*N*=3, 117, 121, 93, 47 and 21 from left to right). (F) Pair-wise scatter plots between different plasticity measurements: modification threshold (θ_m_), percentage weight change at 1 Hz (Δ*w*_1_) and 10 Hz (Δ*w*_10_) and measurements of intrinsic excitability: *R*_in_, *f*_100_, and *f*_150_ for two AMPAR permeability values: 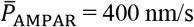 (F) and 600 nm/s (G), across six diameters values (3, 9, 30, 40, 50, 63 μm). The scatter plots are overlaid to corresponding color-coded pair-wise correlation coefficients representing weak pair-wise correlations across diameters and permeability values. Note that θ_m_ did not fall within the tested range of induction frequencies for different models with different diameter values, thus resulting in lesser points for certain diameter values. The axes ranges for each measurement span the entire range of the respective measurements, and are different across different plots.

Considering an example of a single granule cell model, we found that altering the diameter of the neuron had a dramatic impact on the synaptic plasticity profile even when 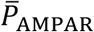 was set at a fixed value (Fig. 7*A–B*). Thus, in the absence of synaptic or intrinsic heterogeneities, structural changes were independently capable of altering synaptic plasticity profiles, introducing a left-ward shift in the plasticity profile with reduction in the diameter (Fig. 7*A–D*). In assessing the impact of synaptic heterogeneities, we plotted modification threshold as a function of diameter for different values of 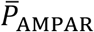 and found the diameter-dependent left-ward shift was observed across 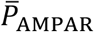 values irrespective of whether the diameter was varied over immature (Fig. 7*C*) and mature (Fig. 7*D*) ranges. As a consequence of leftward shifts in the modification threshold by reduction in diameter, the 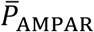 value required for similar modification thresholds was lower in immature neurons (Fig. 7*C*) compared to their developing/mature counterparts (Fig. 7*D*). However, irrespective of the ranges of diameters, increase in 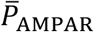 values resulted in an expected leftward shift in the plasticity profiles (Fig. 7*C–D*).

To address the impact of intrinsic heterogeneity on the modification threshold in the context of structural heterogeneity, we chose a specific value of 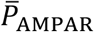 (475 nm/s; Fig. 7*E*) such that there were at least some models with modification threshold value within the tested range of induction frequencies (0.5–25 Hz) across different range of diameter (between 10–63 μm). This was essential because reduction in diameter led to large leftward shifts in the plasticity profile, resulting in a scenario where none of the tested induction frequencies resulted in depression thereby resulting in an indeterminate modification threshold (*e.g.,* diameters 3– 10 μm in Fig. 7*A*). For the fixed value of 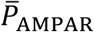, we found that the modification threshold increased as a function of diameter, albeit manifesting considerable heterogeneity in the modification threshold for a given diameter value across different models (Fig. 7*E*). For several models with diameters of 10 μm, 40 μm, 50 μm, and 63 μm, the modification threshold (with 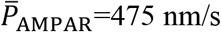) was not within the tested range of induction frequencies (0.5–25 Hz), thus resulting in lesser number of models for those diameters (Fig. 7*E*).

Were there strong relationships between intrinsic and synaptic plasticity measurements across these models across different diameters and different values of 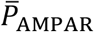? To answer this, we employed three intrinsic measurements (*R*_*in*_, *f*_100_ and *f*_150_) and three measurements of synaptic plasticity (θ_m_, Δ*w*_1_ and Δ*w*_10_), each measured for 6 diameter values (3, 9, 30, 40, 50, 63 μm) and two 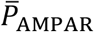 values. We computed Pearson’s correlation coefficients between the intrinsic and synaptic plasticity measurements and found weak pair wise correlations between intrinsic and plasticity measurements across different diameter and permeability values. Together, these analyses demonstrated that immature cells with relatively smaller surface areas showed a lower threshold value for LTP induction, in terms of the induction frequency (Fig. 7*A–B*, Fig. 7*E*) and the baseline synaptic strength (immature, Fig. 7*C* *vs*. mature, Fig. 7*D*). We noted that these observations matched their electrophysiological counterparts showing that immature neurons have lower threshold for plasticity induction compared to mature neurons (Schmidt-Hieber et al., 2004; Ge et al., 2007; Dieni et al., 2013; Aimone et al., 2014).

### Synergistic interactions between different forms of heterogeneities resulted in the emergence of plasticity degeneracy with BCM-like plasticity profiles

At any given time-point, the granule cell population in the DG network comprises of neurons in distinct age groups, spanning the entire range of just-born to fully mature neurons. Thus, based on our analyses so far, the consequent structural and intrinsic heterogeneities could result in distinct plasticity profiles with different ranges of modification thresholds. However, we had demonstrated earlier that similar plasticity profiles could be achieved across different intrinsically heterogeneous GC neurons, if the baseline synaptic strength were adjusted appropriately (Fig. 4). Although intrinsic neural properties and synaptic strength manifested considerable heterogeneities when viewed independently, together they were able to yield very similar plasticity profiles (Fig. 4). Could such plasticity degeneracy manifest even in presence of neurogenesis-induced structural heterogeneity? Could similar plasticity profiles be achieved despite the concomitant expression of intrinsic, synaptic, and structural heterogeneities in the DG neuronal population?

To assess these questions, we first selected six intrinsically distinct GC models (from the population of 126 models) and assigned different values of diameters to each of these six models. We then employed an algorithm to find a synaptic permeability value 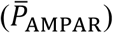 that yielded plasticity profiles endowed with their modification threshold at ~10 Hz with the 900-pulse protocol (Fig. 8*A*). We found the synaptic plasticity profiles for each of these six models, endowed with their respective 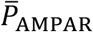 provided by the algorithm, to be similar across the entire range of induction frequencies (0.5–25 Hz) and found them to be very similar (Fig. 8*A*). We then plotted each of the 42 parameters underlying these 6 models (40 intrinsic parameters in Table 1, diameter as the structural parameter, and 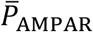 governing the synapse) and found each of them to span their respective ranges (Fig. 8*B*). These analyses illustrate that models built with very different structural, intrinsic, and synaptic properties (Fig. 8*B*) could together yield very similar synaptic plasticity profile (Fig. 8*A*), thus demonstrating the emergence of plasticity degeneracy despite widespread variability in all underlying parameters.

**Figure 8:**
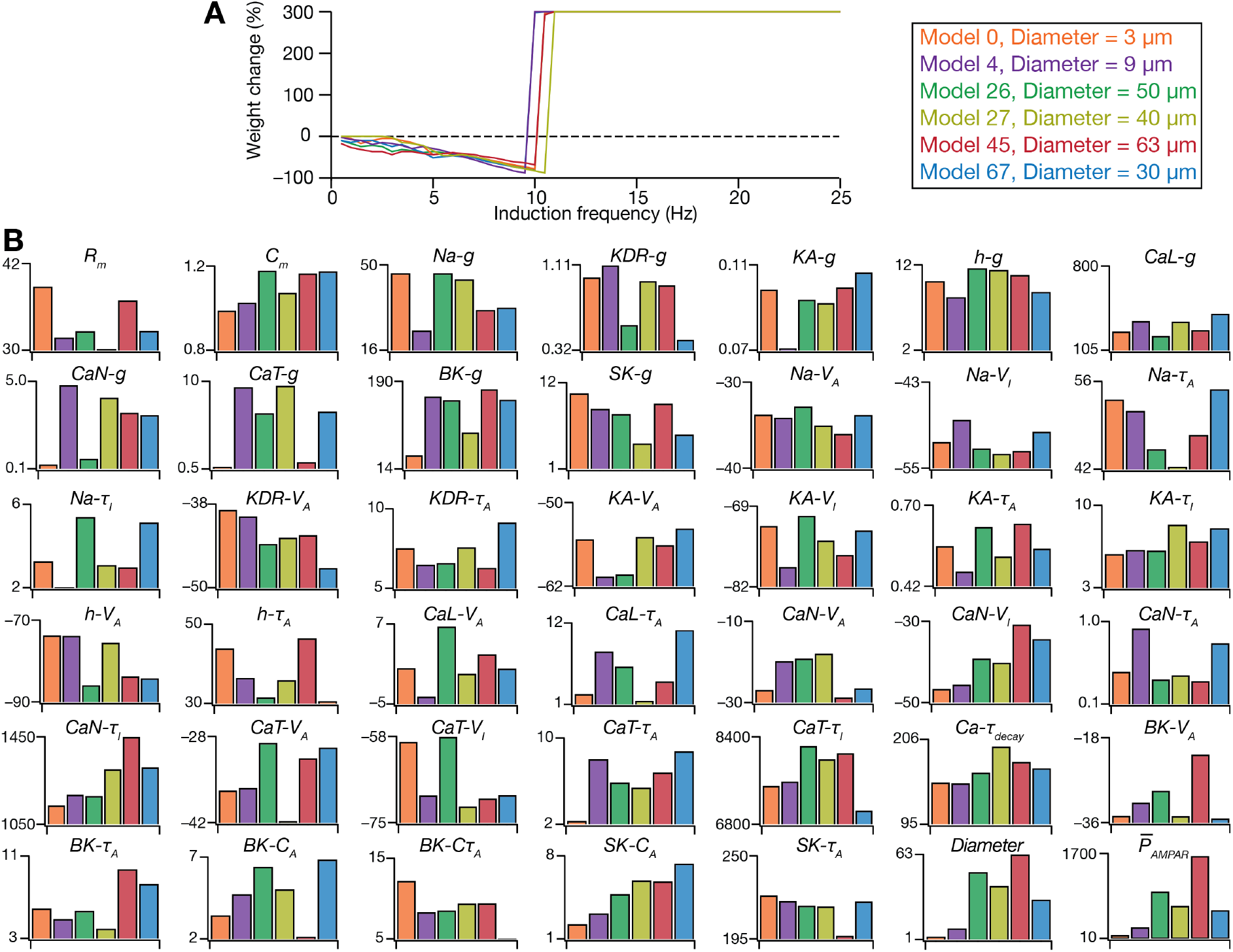
Illustration of degeneracy in the emergence of plasticity profiles spanning biophysical, structural, and synaptic parameters using six models. (A) Frequency-dependent plasticity profiles plotted for six intrinsically disparate models with different diameters and 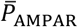 values yield similar plasticity profile with modification threshold at ~10 Hz. (B) Plots, for each of the six models shown in panel *A*, of the 40 intrinsic passive and active properties (listed in Table 1 with units), the diameter (in μm) and the 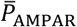 (in nm/s) values required to get the modification threshold to be ~10 Hz. The plots for each of the 40 intrinsic parameters (Table 1) and diameter (1–63 μm) span their entire search range. Note that the ranges of each parameter across the six models is highly variable (B), spanning a large portion of the parameter’s search range, despite the similarity of the plasticity profiles (A).

We expanded the scope of our analyses to span all 126 intrinsically heterogeneous models, each spanning six diameter values (3, 9, 30, 40, 50, and 63 μm) and employed our algorithm to find a 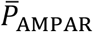 that would yield a modification threshold of ~10 Hz (9.75 ≤ θ_*m*_ ≤ 10.25) in each of these (126 × 6 = 756) models (Fig. 9*A*). We were able to find 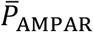 values that yielded similar modification thresholds, with the required 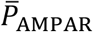 increasing with increase in diameter (Fig. 9*B*). We did not find strong correlations between the 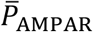 value required for achieving similar plasticity profiles and the respective intrinsic measurements (Fig. 9*C*), thus ruling out the requirement of strong counterbalances between intrinsic and synaptic properties, within each of the 6 assessed diameters. Similarly, there were no strong correlations between the threshold 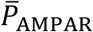 value and each of the 40 intrinsic parameters (Table 1), for models with each of the 6 diameter values (Fig. 9*D*). Together, these results demonstrated that degeneracy in the emergence of plasticity profiles is not dependent on strong pairwise compensations between synaptic properties and individual intrinsic measurements (Fig. 9*C*) or parameters (Fig. 9*D*). These analyses suggest a role for synergistic interactions among structural, intrinsic, and synaptic parameters in yielding *similar* plasticity profiles.

**Figure 9:**
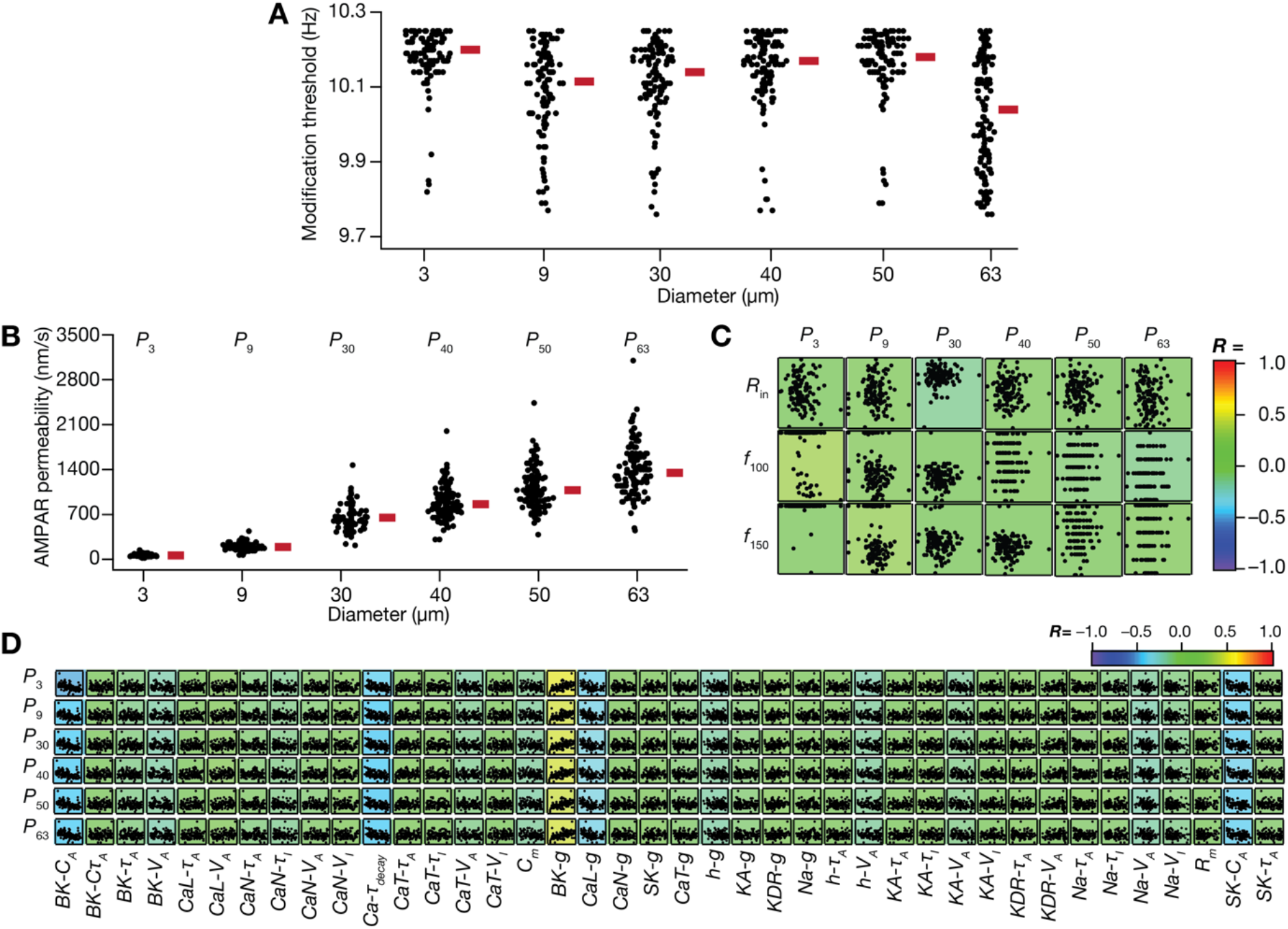
Emergence of plasticity degeneracy due to synergistic interactions between age-dependent structural, synaptic, and intrinsic heterogeneities with weak pair-wise correlations. (A) Plot representing the distribution of modification threshold for all GC models across different diameters to obtain modification threshold of ~10 Hz by adjusting AMPAR permeability for each model. (B) Plot depicting the distribution of AMPAR permeability values required to obtain plasticity profiles with modification threshold of ~10 Hz (shown in panel *A*), across different diameter values. (C) Pair-wise scatter plots between AMPA permeability values depicted in *B* and intrinsic excitability measurements (*R*_in_, *f*_100_, and *f*_150_) across different diameters, overlaid on respective color-coded correlation coefficient values. (D) Pair-wise scatter plots showing distribution of intrinsic parameters across AMPA permeabilities that yielded ~10 Hz modification threshold across different diameters. The scatter plots are overlaid on color-coded pair-wise correlation coefficient values showing weak pair-wise correlations.

### Synergistic interactions between synaptic, intrinsic, and structural heterogeneities governed TBS-induced synaptic plasticity

We repeated our analyses on the impact of the three forms of heterogeneities with the TBS protocol. First, we found that heterogeneities in structural properties could alter the amount of synaptic plasticity achieved with TBS across the intrinsically heterogeneous model population, when structural heterogeneity was introduced by altering diameters to six different values representative of immature and mature granule cell populations. For these analyses, we fixed the 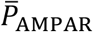 value and found that the amount of plasticity obtained reduced with increasing value of diameter (Fig. 10*A*, left), thus demonstrating a lower threshold on 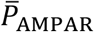 for inducing LTP in immature neurons. We also found that there was no correlation between the amount of plasticity achieved and the respective intrinsic properties, for each value of diameter assessed (Fig. 10*A*, right). Second, to explore plasticity degeneracy with the TBS protocol, we next found 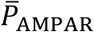 values that yielded similar levels of synaptic plasticity of ~150% (148–152%) for each of the 126 intrinsically heterogeneous models, with 6 different values of diameters (Fig. 10*B–C*). The 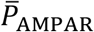 value required for achieving similar plasticity increased with increase in diameter, and did not manifest strong correlations with either the respective intrinsic measurements (Fig. 10*D*) or the intrinsic parameters (Fig. 10*E*) for each value of the diameter.

**Figure 10.**
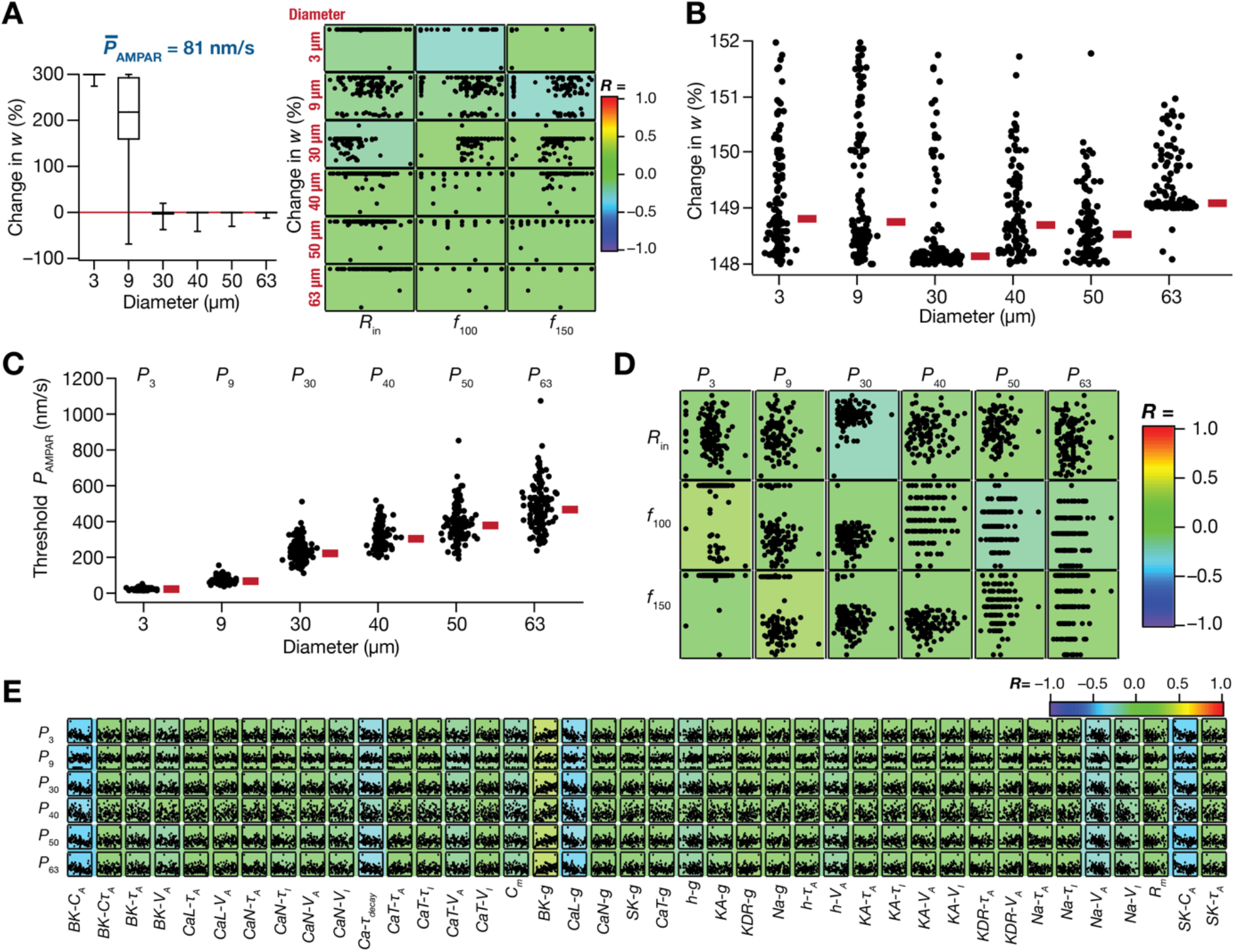
Heterogeneities and degeneracy in synaptic plasticity achieved with theta burst stimulation protocol in models endowed with age-dependent structural, synaptic, and intrinsic heterogeneities. (A) *Left,* Age-dependent structural heterogeneity in the population of GC models translated to heterogeneity in the amount of plasticity achieved with theta burst stimulation (TBS) protocol when baseline synaptic strength was fixed to 81 nm/s. Shown is the amount of plasticity achieved for models in the intrinsically heterogeneous model population, with the diameter altered to assess the impact of structural heterogeneities. *Right*, Pair-wise scatter plots between different plasticity measurements associated with TBS *vs.* measurements of intrinsic excitability: *R*_in_, *f*_100_, and *f*_150_ for a fixed value of baseline synaptic strength, 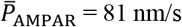, across six diameter values (3, 9, 30, 40, 50, and 63 μm). The scatter plots are overlaid to corresponding color-coded pair-wise correlation coefficients representing weak pair-wise correlations across diameters and permeability values. The axes ranges for each measurement span the entire range of the respective measurements, and are different across different plots. (B–E) Degeneracy in eliciting the same amount of TBS-induced LTP emerges from synergistic interactions between heterogeneities in structural, intrinsic, and synaptic properties. (B) Plot showing that the same amount of LTP (~150%) is obtained in different GC models, across 6 different diameter values to assess the impact of structural heterogeneities, by adjusting 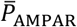. (C) Beeswarm plot representing the range of AMPAR permeabilities required to obtain ~150% LTP shown in *B*, for each of the 6 diameter values. (D) Pair-wise scatter plots between permeability parameter in *C* and three intrinsic measurements of the respective models, overlaid on the corresponding color-coded correlation coefficients. Plots are shown for each of the 6 diameter values. (E) Pair-wise scatter plots between permeability parameter in *B* and intrinsic parameters spanning all GC models. Overlaid are respective color-coded correlation values.

Together, our analyses demonstrated that each of intrinsic (Fig. 2*D–F*, Fig. 5*D*, Fig. 7*E*, Fig. 10*A*), synaptic (Fig. 2*D–F*, Fig. 5*C–D*, Fig. 7*C–D*), and structural (Fig. 7*E*, Fig. 10*A*) heterogeneities could independently introduce heterogeneities in the plasticity profiles, irrespective of the protocol employed. However, when they coexpress, these disparate forms of heterogeneities could synergistically interact with each other to yield very similar plasticity profiles (Fig. 4*A–C*, Fig. 6*A–B*, Fig. 8, Fig 9*A–B*, Fig. 10*B–C*), irrespective of the induction protocol employed. Across our analyses spanning different plasticity protocols, assessing heterogeneities or degeneracy in plasticity profiles, we did not find strong correlations between synaptic properties plotted against intrinsic measurements (Fig. 3, Fig. 4*D*, Fig. 5*E*, Fig. 6*C*, Fig. 7*F–G*, Fig. 9*C*, Fig. 10*D*) or intrinsic parameters (Fig. 4*E*, Fig. 6*D*, Fig. 9*D*, Fig. 10*E*) of the model populations. These results suggested that the measurements of intrinsic excitability and temporal summation are not sufficiently strong to impose specific synaptic plasticity profiles.

### Importance of adult neurogenesis-induced structural heterogeneities in lowering plasticity induction threshold and recruiting engram cells based on intrinsic excitability

In our analyses thus far, we noted that intrinsic excitability parameters were not strong enough to constrain synaptic plasticity induction, with a consistent lack of strong correlations between synaptic plasticity measurements and intrinsic excitability (Figs. 4–7, Figs. 9–10). This is in contrast with the literature where a critical role for intrinsic excitability has been postulated in reducing the threshold for plasticity induction and in individual neurons being recruited as engram cells for a new context (Schmidt-Hieber et al., 2004; Ge et al., 2007; Yiu et al., 2014; Park et al., 2016; Josselyn and Frankland, 2018; Lodge and Bischofberger, 2019; Pignatelli et al., 2019; Josselyn and Tonegawa, 2020; Lau et al., 2020; Huckleberry and Shansky, 2021). How do we reconcile these observations? Thus far in our correlation analyses, we have focused *independently* on mature *vs.* immature populations, treating analyses within each diameter to be independent of others (Figs. 7, 9–10). However, in physiological scenarios where there is coexistence of cells of different ages, it is important to ask if immature cells have an *advantage* over their mature counterparts in manifesting lower threshold for plasticity, and thereby being recruited by afferent inputs. It is therefore essential that the analyses *span all ages of cells* rather than treating them as independent populations.

Therefore, we plotted the amount of plasticity induced by the 900-pulse protocol at *f*_*i*_=1 Hz (Δ*w*_1_; Fig. 11*A–B*) and the modification threshold obtained with the 900-pulse protocol (θ_m_; Fig. 11*C–D*), computed with a fixed value of 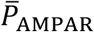, against respective intrinsic excitability measurements (*R*_*in*_ and *f*_100_) spanning *all diameter values* across all neurons in the intrinsically heterogeneous population. We found strong relationships of synaptic plasticity measurements with measurements of intrinsic excitability (Fig. 11*A–D*). Specifically, our analyses showed that the amount of induced plasticity was higher in neurons with high excitability (Fig. 11*A–B*) and that the modification threshold manifested a strong leftward shift with increased excitability (Fig. 11*C–D*). Thus, while intrinsic excitability was insufficient to impose strong correlations on synaptic plasticity measurements in the absence of structural heterogeneities in the neural population (Figs. 7, 9–10), quantitative introduction of structural heterogeneities allows the consequent intrinsic excitability changes to impose strong constraints on the synaptic plasticity profiles. In other words, although DG granule cells are endowed with considerable baseline heterogeneities, the introduction of an immature population through adult neurogenesis is essential for these neurons to be specifically recruited in a new context during engram formation (Schmidt-Hieber et al., 2004; Yiu et al., 2014; Park et al., 2016; Josselyn and Frankland, 2018; Lodge and Bischofberger, 2019; Pignatelli et al., 2019; Josselyn and Tonegawa, 2020; Lau et al., 2020; Huckleberry and Shansky, 2021).

**Figure 11.**
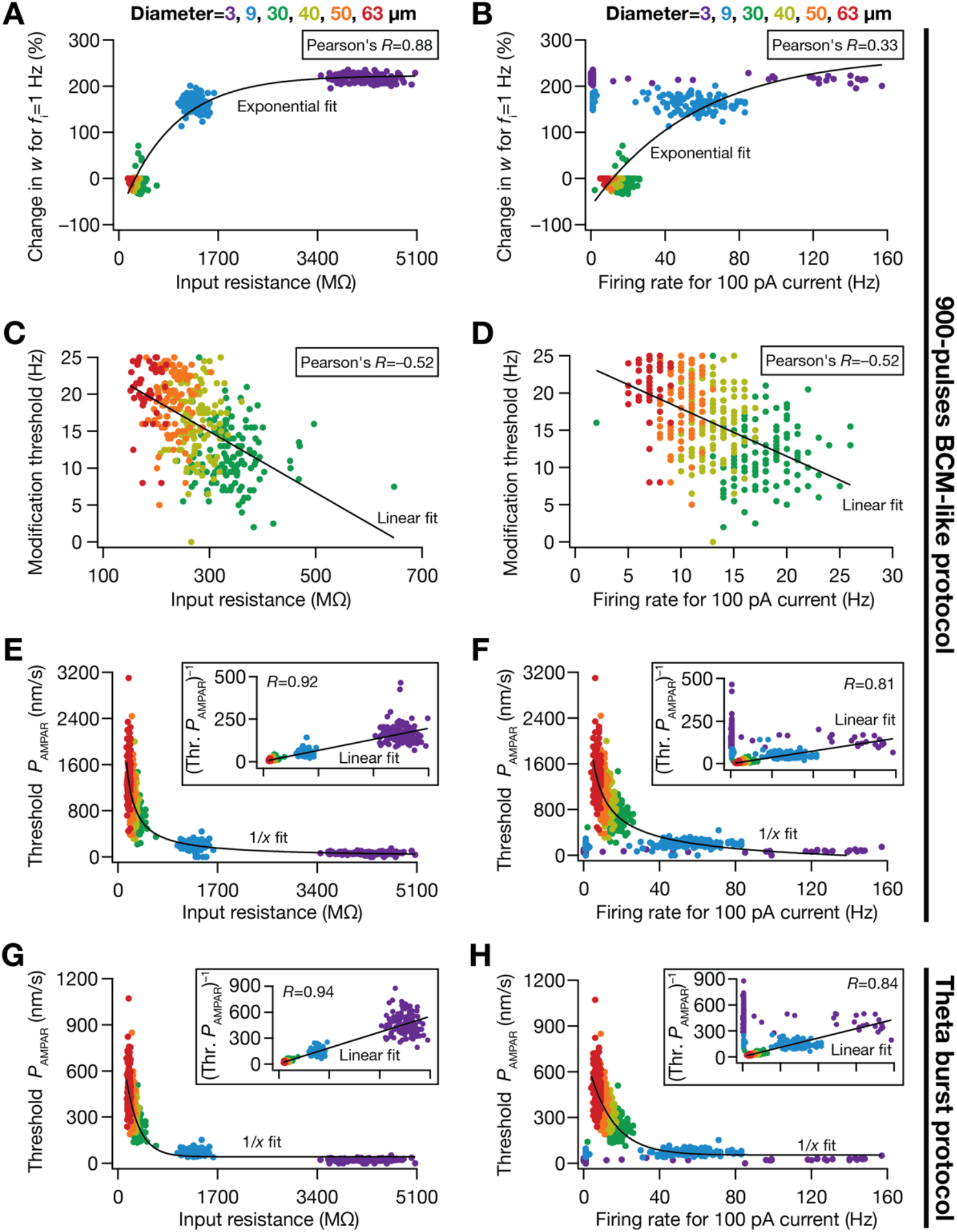
The dominant role of structural heterogeneities in regulating plasticity profiles with the BCM-like and TBS plasticity protocols. (A–B) Percentage weight change at 1 Hz (Δ*w*_1_) with the 900-pulses protocol plotted against input resistance (A) and firing rate for 100 pA current injection (B) for models in the intrinsically heterogeneous population, with each model assessed at 6 different diameter values (3, 9, 30, 40, 50, and 63 μm). (C–D) Same as (A–B), plotted for modification threshold (θ_m_) on the *y* axis. For panels (A–D), the data from Fig. 7G 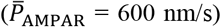 is plotted together for all diameters. (E–F) Same as (A–B), plotted for the AMPAR permeabilities required to achieve a modification threshold of ~10 Hz (with the 900-pulses protocol), referred to as Threshold *P*_AMPAR_, on the *y* axis. For panels (E–F), the data from Fig. 9*C* is plotted together for all diameters. The insets in panels (E) and (F) depict the inverse of threshold *P*_AMPAR_ plotted against *R*_in_ or *f*_100_, respectively, to illustrate the 1/*x* relationship between Threshold *P*_AMPAR_ *vs*. *R*_+3_ and Threshold *P*_AMPAR_ *vs*. *f*_100_. (G–H) Same as (E–F), but plotted Threshold *P*_AMPAR_ was computed to achieve ~150% synaptic plasticity with the TBS protocol. For panels (G–H), the data from Fig. 10*D* is plotted together for all diameters.

There are lines of evidence that the synaptic strength of inputs to immature DG granule cells is lower compared to their mature counterparts (Mongiat et al., 2009; Dieni et al., 2016; Li et al., 2017). Are immature neurons with such low baseline synaptic strengths capable of effectuating synaptic plasticity comparable to their mature counterparts? To address this, we first plotted the threshold *P*_*AMPAR*_ required to elicit the same modification threshold (of ~10 Hz) with the 900-pulses protocol as functions of intrinsic excitability measurements, spanning *all diameter* values across all neurons in the intrinsically heterogeneous population (Fig. 11*C–D*). We found that in immature neurons with high excitability, even small values of *P*_*AMPAR*_ were sufficient to achieve synaptic plasticity profiles comparable with their mature counterparts (Fig. 11*E*). Importantly, although threshold *P*_*AMPAR*_ values from *individual* neuronal populations of different diameters did not manifest strong correlations with intrinsic measurements (Fig. 9*C*), there was a strong inverse relationship between threshold *P*_*AMPAR*_ values and *R*_*in*_ (Fig. 11*E*) as well as *f*_100_ (Fig. 11*F*) when neurons with all diameters were considered together. Finally, we found that small values of *P*_*AMPAR*_ were sufficient to achieve synaptic plasticity of ~150% with the TBS protocol in immature neurons (Fig. 11*G*). Strong inverse relationships manifested between threshold *P*_*AMPAR*_ and measurements of intrinsic excitability with the TBS protocol as well (Fig. 11*G–H*). Together, these analyses demonstrated the essential requirement of structural heterogeneity comprising immature neuronal populations in specifically recruiting high-excitability populations to encode specific contexts during engram cell formation, and in ensuring that sufficient plasticity is induced despite low density of synaptic receptors in immature neurons.

## DISCUSSION

### Heterogeneities spawn heterogeneities: Disparate forms of biophysical and structural heterogeneities could independently drive physiologically crucial heterogeneities in plasticity profiles

We constructed multiple populations of dentate gyrus granule cell models to reflect heterogeneities in neuronal passive properties, ion-channel properties, calcium-handling mechanisms, synaptic strength, and neural structure of DG granule cells of different ages. Each of these heterogeneities was incorporated into our model populations with strong physiological constraints on multiple intrinsic properties (Table 2), thus ensuring the physiological relevance of our conclusions. We employed two well-established synaptic plasticity protocols to demonstrate that each of intrinsic, synaptic, and structural heterogeneities independently result in heterogeneities in the amount of plasticity induced. These observations held for both plasticity protocols, one involving 900 pulses of different induction frequencies and another employing theta-burst stimulation. In electrophysiological experiments assessing synaptic plasticity, there is pronounced heterogeneity in the amount of plasticity induced with any induction protocol. Specifically, whereas the same protocol might elicit 300% LTP in certain neurons, in other neurons of the same subtype in the same brain region, the protocol results in 10% LTP. Such neuron-to-neuron and animal-to-animal variability in the amount of plasticity induced is typically not analyzed quantitatively, with the data typically represented using summary statistics and interpretations drawn from the average plasticity across different cells from different animals. However, given the role of such differential plasticity across different neurons in resource allocation and in engram formation, it is critical not just to report these heterogeneities but also to assess the mechanisms underlying such cell-to-cell differences.

To emphasize the critical roles played by these plasticity heterogeneities across different cells and different synapses, let us consider an extreme scenario where these heterogeneities were absent. This would translate to all synapses across all cells undergoing the *same* amount of plasticity for any given context, together resulting in the absence of context-dependent recruitment/allocation of synapses or cells that is critical for engram cell formation and decorrelation. From the engram cell formation perspective, there are several lines of evidence to suggest context-dependent plasticity in a *subset* of cells that are recruited to encode a new context (Schmidt-Hieber et al., 2004; Yiu et al., 2014; Park et al., 2016; Josselyn and Frankland, 2018; Lodge and Bischofberger, 2019; Pignatelli et al., 2019; Josselyn and Tonegawa, 2020; Lau et al., 2020). In addition, afferent connectivity has been demonstrated to be a dominant mediator of neural decorrelation (Mishra and Narayanan, 2019), and there are strong lines of evidence that afferent connectivity is actively driven by differences in plasticity profiles across different granule cells (Schmidt-Hieber et al., 2004; Aimone et al., 2006; Ge et al., 2007; Aimone et al., 2009; Aimone et al., 2014; Li et al., 2017; Lodge and Bischofberger, 2019; Luna et al., 2019). Thus, in the absence of plasticity heterogeneities where similar plasticity manifest across all neurons, the critical role of differential plasticity in mediating differential connectivity to neurons in the DG during encoding and storage process would be hampered. Our study explores the mechanistic basis for such heterogeneity, and traces the potential origins to the pronounced heterogeneities in intrinsic, synaptic, and structural properties of DG granule cells. These analyses and the important physiological roles played by differential plasticity in encoding processes together emphasize the need for studies that assess neural plasticity to quantitatively report plasticity heterogeneities and to trace their origins, under physiological or pathological conditions.

### Heterogeneities underlying degeneracy: Synergistic interactions among different forms of biophysical and structural heterogeneities could yield similar plasticity profiles

We demonstrated that even the expression of heterogeneities in *all of* structural, synaptic, and intrinsic neuronal properties does not necessarily have to translate to heterogeneities in synaptic plasticity profiles. Specifically, we showed that very similar plasticity profiles could be achieved with disparate combinations of neuronal passive properties, ion-channel properties, calcium-handling mechanisms, synaptic strength, and neural structure of DG granule cells of different ages (Fig. 4, Fig. 6, Figs. 8–10). Independently observed, these properties manifested widespread heterogeneities with no pair-wise relationships (Fig. 4, Fig. 6, Figs. 9–10). But when seen together, these heterogeneities synergistically interacted with each other to achieve the functional goal of degeneracy in the synaptic plasticity profiles. There are computational (Beining et al., 2017; Mishra and Narayanan, 2019, 2021c) and electrophysiological (Mishra and Narayanan, 2021a) lines of evidence for DG granule cells to manifest ion-channel degeneracy in the expression of their characteristic intrinsic properties. However, similarity in baseline neuronal properties of different neurons does not necessarily translate to similarity in terms of how these neurons react to plasticity-inducing stimuli (Anirudhan and Narayanan, 2015; Srikanth and Narayanan, 2015; Rathour and Narayanan, 2019).

Here, we have demonstrated and expanded the scope for the expression of degeneracy in DG granule cells beyond ion-channel degeneracy, and beyond achieving characteristic intrinsic properties. Specifically, we have demonstrated the manifestation of degeneracy in the emergence of plasticity profiles, independently for two different induction protocols, with the analyses *concomitantly* incorporating structural heterogeneities driven by adult-neurogenesis, heterogeneities in intrinsic neuronal properties (spanning passive electrical properties, ion-channel properties, and calcium handling mechanisms), and heterogeneities in synaptic strength (Figs. 8–10). Importantly, this form of degeneracy was demonstrated in a heterogeneous population of neurons that manifested physiologically constrained (Table 2) neural properties, including those which were not initially assessed (Fig. 1), and which also were physiologically matched with neural excitability measurements of immature cells by changes in surface area. Thus, this population of neurons manifested degeneracy in the expression of physiologically matched neural intrinsic properties and showed plasticity degeneracy with the concomitant expression of all forms of neural heterogeneities. In comparing with previous studies on degeneracy, we note that these studies accounted for degeneracy either in characteristic neuronal intrinsic properties (Mishra and Narayanan, 2019, 2021c, a) or in plasticity profiles (Anirudhan and Narayanan, 2015), but not together. Our study demonstrates the expression of degeneracy in the concomitant emergence of characteristic neuronal intrinsic properties and of characteristic plasticity profiles, while considering a superset of model parameters and measurements that span all ages of granule cells in the dentate gyrus.

The predominant implication for the expression of degeneracy in the concomitant emergence of intrinsic properties and plasticity profiles is the explosion in the degrees of freedom available for the neurons to achieve these characteristic features, thereby providing multiple routes to achieving functional robustness (Edelman and Gally, 2001; Rathour and Narayanan, 2019; Goaillard and Marder, 2021). In addition, given the expression of such degeneracy, it is essential that the theoretical and experimental analyses recognize that the mappings between structural components and functional outcomes are many-to-many, and avoid reductionist oversimplifications of structure-function relationships (Rathour and Narayanan, 2019; Goaillard and Marder, 2021; Mishra and Narayanan, 2021a, c). An essential first step in achieving this is the quantification of heterogeneities in the dependencies of each of the several functional outcomes on the individual structural components (Mishra and Narayanan, 2021a, c) — with examples including the different ion-channel/receptor subtypes and the distinct calcium handling mechanisms. It is therefore critical that experimental as well as computational analyses explicitly account for heterogeneities in neural circuit properties and for the expression of degeneracy in the emergence of baseline neural properties and plasticity profiles.

### Dominant role of structural heterogeneities in introducing plasticity heterogeneities across neurons: Selective recruitment and resource allocation during engram cell formation

A role for intrinsic excitability has been postulated in reducing the threshold for plasticity induction and in individual neurons being recruited as engram cells, towards encoding a new context (Schmidt-Hieber et al., 2004; Ge et al., 2007; Yiu et al., 2014; Park et al., 2016; Josselyn and Frankland, 2018; Lodge and Bischofberger, 2019; Pignatelli et al., 2019; Josselyn and Tonegawa, 2020; Lau et al., 2020). However, in the individual population of mature or immature cells, we demonstrated that intrinsic excitability and temporal summation heterogeneities were insufficient to impose strong constraints on plasticity-related measurements (Fig. 3, Fig. 4*D*, Fig. 5*E*, Fig. 6*C*, Fig. 7*F–G*, Fig. 9*C*, Fig. 10*D*). However, when the entire population covering mature and immature cells were considered *together*, it was clear that there were strong relationships between intrinsic excitability and measurements associated with synaptic plasticity (Fig. 11). These results highlighted the dominance of structural heterogeneities, introduced through adult neurogenesis, in introducing heterogeneities in plasticity profiles (discussed above) that are essential for effective execution of encoding and storage roles of the dentate gyrus.

We explored a range of AMPAR strengths across neurons with different diameters and demonstrate that similar levels of synaptic plasticity could be achieved despite the low synaptic strength (Fig. 11*E–H*) that is observed onto these immature neurons (Mongiat et al., 2009; Dieni et al., 2016; Li et al., 2017). The lower ranges of AMPAR strengths may indeed be an essential requirement for keeping the plasticity in a useful physiological range, because higher AMPAR strengths in immature neurons might result in large magnitude and unstable plasticity dynamics. Together, the amplified heterogeneities that emerge as a consequence of the manifestation of adult neurogenesis form a crucial lynchpin in defining plasticity heterogeneity across different neurons and in defining a role for intrinsic excitability in recruitment/allocation of engram cells in the dentate gyrus. Specifically, the heterogeneities typically seen in mature cells (with input resistance in the range of 100–300 MΩ) are not sufficient to enforce strong correlations of neural excitability measurements with the expression of synaptic plasticity. Higher input resistance values (in the GΩ range) that distinguish the immature neurons from their mature counterparts are essential to significantly lower the synaptic plasticity induction threshold and drive the recruitment of these as engram cells for new contexts.

### Future directions

Although our model population spanned all forms of heterogeneities and was physiologically constrained in several ways, there are some limitations in the model and in parametric choices. First, the computational cost for each plasticity simulation involved either 900 pulses of different *f*_*i*_ values ranging from 0.5–25 Hz in steps of 0.5 Hz, or TBS repeated for 150 times, with these repeated for each of the 126 intrinsically heterogeneous models with different 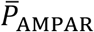 and several diameter values. To partially offset for this tremendous computational cost, we had employed a single-compartmental model to assess the impact of neural-circuit heterogeneities on plasticity profiles. However, it is essential that future studies account for morphological reconstructions of DG granule cells with experimentally determined somato-dendritic distributions of channels and receptors and assess plasticity profiles for synapses placed at different somato-dendritic locations (Sjostrom and Hausser, 2006). These studies could also specifically employ immature *vs.* mature dendritic morphologies rather than introducing surface area changes through change in diameter. Such analyses would provide insights about the impact of morphological heterogeneities, spanning baseline *vs*. adult-neurogenesis driven structural differences in the dendritic arbor, on location-dependent plasticity profiles spanning the stratified synaptic inputs from lateral *vs*. medial entorhinal cortices.

Second, as our focus here was on excitatory synaptic plasticity, we have not incorporated inhibition into our analyses. However, given the DG circuitry that recruits a diverse set of interneurons that impinge along different locations of the somato-dendritic arbor (Freund and Buzsaki, 1996; Andersen et al., 2006; Amaral et al., 2007; Houser, 2007; Dieni et al., 2013; Elgueta and Bartos, 2019), it is essential that the impact of heterogeneities in inhibitory synaptic inputs on plasticity profiles are also assessed in more detail. In this context, there are lines of evidence that the inhibitory neurotransmitter GABA exerts functionally critical excitatory influences on the immature cells, and through the process of maturation shifts to being inhibitory (Ge et al., 2006; Chancey et al., 2013; Heigele et al., 2016). Thus, future studies that account for inhibition should also assess the impact of this switch in GABAergic impact on immature *vs.* mature granule cells and their plasticity profiles.

Third, whereas our *cellular-scale* analysis has focused on the biophysical and structural heterogeneities as sources of plasticity heterogeneities, there are other potential sources for plasticity heterogeneities. At the molecular scale, it is possible that heterogeneities in the expression of plasticity related molecules (and associated signaling cascades) across synapses and across neurons of the same subtype could mediate plasticity heterogeneities. At the network scale, when multiple neurons are considered, pre-existing afferent as well as local connectivity onto these neurons could form yet another potential source of plasticity heterogeneities. Thus, future studies could expand the analyses of plasticity heterogeneity beyond the cellular scale to encompass network- and molecular-scale components that could drive plasticity heterogeneities. Finally, our analyses here was limited to synaptic plasticity. However, plasticity in the DG granule cells is known to span synaptic and intrinsic properties (Bliss and Lomo, 1973; Lopez-Rojas et al., 2016; Mishra and Narayanan, 2020b, 2021b), thus necessitating the need for future studies to assess the impact of neural heterogeneities on *conjunctive* intrinsic and synaptic plasticity.

## Acknowledgments

The authors thank the members of the cellular neurophysiology laboratory for helpful comments on a draft version of this manuscript. This work was supported by the Wellcome Trust-DBT India Alliance (Senior fellowship to RN; IA/S/16/2/502727), Human Frontier Science Program (HFSP) Organization (RN), the Department of Biotechnology through the DBT-IISc partnership program (RN), the Revati & Satya Nadham Atluri Chair at IISc (RN), the Department of Science and Technology (RN), and the Ministry of Human Resource Development (RN & PM).

## REFERENCES

Aimone JB, Wiles J, Gage FH (2006) Potential role for adult neurogenesis in the encoding of time in new memories. Nat Neurosci 9:723–727.

Aimone JB, Wiles J, Gage FH (2009) Computational influence of adult neurogenesis on memory encoding. Neuron 61:187–202.

Aimone JB, Li Y, Lee SW, Clemenson GD, Deng W, Gage FH (2014) Regulation and function of adult neurogenesis: from genes to cognition. Physiol Rev 94:991–1026.

Amaral DG, Scharfman HE, Lavenex P (2007) The dentate gyrus: fundamental neuroanatomical organization (dentate gyrus for dummies). Prog Brain Res 163:3–22.

Andersen P, Morris R, Amaral D, Bliss T, O’Keefe J (2006) The hippocampus book. New York, USA: Oxford University Press.

Anirudhan A, Narayanan R (2015) Analogous synaptic plasticity profiles emerge from disparate channel combinations. J Neurosci 35:4691–4705.

Aradi I, Holmes WR (1999) Role of multiple calcium and calcium-dependent conductances in regulation of hippocampal dentate granule cell excitability. J Comput Neurosci 6:215–235.

Ashhad S, Narayanan R (2013) Quantitative interactions between the A-type K+ current and inositol trisphosphate receptors regulate intraneuronal Ca2+ waves and synaptic plasticity. J Physiol 591:1645–1669.

Beck H, Goussakov IV, Lie A, Helmstaedter C, Elger CE (2000) Synaptic plasticity in the human dentate gyrus. J Neurosci 20:7080–7086.

Beining M, Mongiat LA, Schwarzacher SW, Cuntz H, Jedlicka P (2017) T2N as a new tool for robust electrophysiological modeling demonstrated for mature and adult-born dentate granule cells. eLife 6.

Bienenstock EL, Cooper LN, Munro PW (1982) Theory for the development of neuron selectivity: orientation specificity and binocular interaction in visual cortex. J Neurosci 2:32–48.

Bliss TV, Lomo T (1973) Long-lasting potentiation of synaptic transmission in the dentate area of the anaesthetized rabbit following stimulation of the perforant path. J Physiol 232:331–356.

Bliss TV, Gardner-Medwin AR (1973) Long-lasting potentiation of synaptic transmission in the dentate area of the unanaestetized rabbit following stimulation of the perforant path. J Physiol 232:357–374.

Canavier CC (1999) Sodium dynamics underlying burst firing and putative mechanisms for the regulation of the firing pattern in midbrain dopamine neurons: a computational approach. J Comput Neurosci 6:49–69.

Carnevale TN, Hines LM (2006) The NEURON Book. United Kingdom: Cambridge University Press.

Chancey JH, Adlaf EW, Sapp MC, Pugh PC, Wadiche JI, Overstreet-Wadiche LS (2013) GABA depolarization is required for experience-dependent synapse unsilencing in adult-born neurons. J Neurosci 33:6614–6622.

Chen C (2004) ZD7288 inhibits postsynaptic glutamate receptor-mediated responses at hippocampal perforant path-granule cell synapses. The European journal of neuroscience 19:643–649.

Cooper LN, Bear MF (2012) The BCM theory of synapse modification at 30: interaction of theory with experiment. Nature reviews Neuroscience 13:798–810.

Davis CD, Jones FL, Derrick BE (2004) Novel environments enhance the induction and maintenance of long-term potentiation in the dentate gyrus. J Neurosci 24:6497–6506.

Destexhe A, Babloyantz A, Sejnowski TJ (1993) Ionic mechanisms for intrinsic slow oscillations in thalamic relay neurons. Biophysical journal 65:1538–1552.

Dieni CV, Nietz AK, Panichi R, Wadiche JI, Overstreet-Wadiche L (2013) Distinct determinants of sparse activation during granule cell maturation. J Neurosci 33:19131–19142.

Dieni CV, Panichi R, Aimone JB, Kuo CT, Wadiche JI, Overstreet-Wadiche L (2016) Low excitatory innervation balances high intrinsic excitability of immature dentate neurons. Nature communications 7:11313.

Dudek SM, Bear MF (1992) Homosynaptic long-term depression in area CA1 of hippocampus and effects of N-methyl-D-aspartate receptor blockade. Proc Natl Acad Sci U S A 89:4363–4367.

Edelman GM, Gally JA (2001) Degeneracy and complexity in biological systems. Proc Natl Acad Sci U S A 98:13763–13768.

Elgueta C, Bartos M (2019) Dendritic inhibition differentially regulates excitability of dentate gyrus parvalbumin-expressing interneurons and granule cells. Nature communications 10:5561.

Evans JD (1996) Straightforward statistics for the behavioral sciences. Boston, MA, USA: Brooks/Cole Pub Co.

Freund TF, Buzsaki G (1996) Interneurons of the hippocampus. Hippocampus 6:347–470.

Ge S, Yang CH, Hsu KS, Ming GL, Song H (2007) A critical period for enhanced synaptic plasticity in newly generated neurons of the adult brain. Neuron 54:559–566.

Ge S, Goh EL, Sailor KA, Kitabatake Y, Ming GL, Song H (2006) GABA regulates synaptic integration of newly generated neurons in the adult brain. Nature 439:589–593.

Goaillard JM, Marder E (2021) Ion Channel Degeneracy, Variability, and Covariation in Neuron and Circuit Resilience. Annu Rev Neurosci 44:335–357.

Goldman DE (1943) Potential, Impedance, and Rectification in Membranes. The Journal of general physiology 27:37–60.

Greenstein YJ, Pavlides C, Winson J (1988) Long-term potentiation in the dentate gyrus is preferentially induced at theta rhythm periodicity. Brain Res 438:331–334.

Heigele S, Sultan S, Toni N, Bischofberger J (2016) Bidirectional GABAergic control of action potential firing in newborn hippocampal granule cells. Nat Neurosci 19:263–270.

Hodgkin AL, Katz B (1949) The effect of sodium ions on the electrical activity of giant axon of the squid. J Physiol 108:37–77.

Hodgkin AL, Huxley AF (1952) A quantitative description of membrane current and its application to conduction and excitation in nerve. J Physiol 117:500–544.

Honnuraiah S, Narayanan R (2013) A calcium-dependent plasticity rule for HCN channels maintains activity homeostasis and stable synaptic learning. PLoS One 8:e55590.

Houser CR (2007) Interneurons of the dentate gyrus: an overview of cell types, terminal fields and neurochemical identity. Prog Brain Res 163:217–232.

Huckleberry KA, Shansky RM (2021) The unique plasticity of hippocampal adult-born neurons: Contributing to a heterogeneous dentate. Hippocampus 31:543–556.

Jahr CE, Stevens CF (1990) Voltage dependence of NMDA-activated macroscopic conductances predicted by single-channel kinetics. J Neurosci 10:3178–3182.

Johnston D, Christie BR, Frick A, Gray R, Hoffman DA, Schexnayder LK, Watanabe S, Yuan LL (2003) Active dendrites, potassium channels and synaptic plasticity. Philos Trans R Soc Lond B Biol Sci 358:667–674.

Josselyn SA, Frankland PW (2018) Memory Allocation: Mechanisms and Function. Annu Rev Neurosci 41:389–413.

Josselyn SA, Tonegawa S (2020) Memory engrams: Recalling the past and imagining the future. Science 367:eaaw4325.

Kobayashi K, Iwai T, Sasaki-Hamada S, Kamanaka G, Oka J (2013) Exendin (5-39), an antagonist of GLP-1 receptor, modulates synaptic transmission via glutamate uptake in the dentate gyrus. Brain Res 1505:1–10.

Koranda JL, Masino SA, Blaise JH (2008) Bidirectional synaptic plasticity in the dentate gyrus of the awake freely behaving mouse. J Neurosci Methods 167:160–166.

Krueppel R, Remy S, Beck H (2011) Dendritic integration in hippocampal dentate granule cells. Neuron 71:512–528.

Larson J, Munkacsy E (2015) Theta-burst LTP. Brain Res 1621:38–50.

Lau JMH, Rashid AJ, Jacob AD, Frankland PW, Schacter DL, Josselyn SA (2020) The role of neuronal excitability, allocation to an engram and memory linking in the behavioral generation of a false memory in mice. Neurobiology of learning and memory 174:107284.

Li L, Sultan S, Heigele S, Schmidt-Salzmann C, Toni N, Bischofberger J (2017) Silent synapses generate sparse and orthogonal action potential firing in adult-born hippocampal granule cells. eLife 6.

Lodge M, Bischofberger J (2019) Synaptic properties of newly generated granule cells support sparse coding in the adult hippocampus. Behav Brain Res 372:112036.

Lopez-Rojas J, Heine M, Kreutz MR (2016) Plasticity of intrinsic excitability in mature granule cells of the dentate gyrus. Scientific reports 6:21615.

Lubke J, Frotscher M, Spruston N (1998) Specialized electrophysiological properties of anatomically identified neurons in the hilar region of the rat fascia dentata. Journal of neurophysiology 79:1518–1534.

Luna VM, Anacker C, Burghardt NS, Khandaker H, Andreu V, Millette A, Leary P, Ravenelle R, Jimenez JC, Mastrodonato A, Denny CA, Fenton AA, Scharfman HE, Hen R (2019) Adult-born hippocampal neurons bidirectionally modulate entorhinal inputs into the dentate gyrus. Science 364:578–583.

Marder E, Taylor AL (2011) Multiple models to capture the variability in biological neurons and networks. Nat Neurosci 14:133–138.

Mayer ML, Westbrook GL (1987) Permeation and block of N-methyl-D-aspartic acid receptor channels by divalent cations in mouse cultured central neurones. J Physiol 394:501–527.

McHugh TJ, Jones MW, Quinn JJ, Balthasar N, Coppari R, Elmquist JK, Lowell BB, Fanselow MS, Wilson MA, Tonegawa S (2007) Dentate gyrus NMDA receptors mediate rapid pattern separation in the hippocampal network. Science 317:94–99.

Mishra P, Narayanan R (2019) Disparate forms of heterogeneities and interactions among them drive channel decorrelation in the dentate gyrus: Degeneracy and dominance. Hippocampus 29:378–403.

Mishra P, Narayanan R (2020a) Heterogeneities in intrinsic excitability and frequency-dependent response properties of granule cells across the blades of the rat dentate gyrus. Journal of neurophysiology 123:755–772.

Mishra P, Narayanan R (2020b) Plasticity manifolds: Conjunctive changes in multiple ion channels mediate activity-dependent plasticity in hippocampal granule cells. bioRxiv http://doi.org/10.1101/747550.

Mishra P, Narayanan R (2021a) Ion-channel degeneracy: Multiple ion channels heterogeneously regulate intrinsic physiology of rat hippocampal granule cells. Physiol Rep 9:e14963.

Mishra P, Narayanan R (2021b) Stable continual learning through structured multiscale plasticity manifolds. Current opinion in neurobiology 70:51–63.

Mishra P, Narayanan R (2021c) Ion-channel regulation of response decorrelation in a heterogeneous multi-scale model of the dentate gyrus. Curr Res Neurobiol 2:100007.

Mongiat LA, Esposito MS, Lombardi G, Schinder AF (2009) Reliable activation of immature neurons in the adult hippocampus. PloS one 4:e5320.

Narayanan R, Johnston D (2008) The h channel mediates location dependence and plasticity of intrinsic phase response in rat hippocampal neurons. J Neurosci 28:5846–5860.

Narayanan R, Johnston D (2010) The h current is a candidate mechanism for regulating the sliding modification threshold in a BCM-like synaptic learning rule. Journal of neurophysiology 104:1020–1033.

Nolan MF, Malleret G, Dudman JT, Buhl DL, Santoro B, Gibbs E, Vronskaya S, Buzsaki G, Siegelbaum SA, Kandel ER, Morozov A (2004) A behavioral role for dendritic integration: HCN1 channels constrain spatial memory and plasticity at inputs to distal dendrites of CA1 pyramidal neurons. Cell 119:719–732.

Overstreet-Wadiche LS, Bensen AL, Westbrook GL (2006a) Delayed development of adult-generated granule cells in dentate gyrus. J Neurosci 26:2326–2334.

Overstreet-Wadiche LS, Bromberg DA, Bensen AL, Westbrook GL (2006b) Seizures accelerate functional integration of adult-generated granule cells. J Neurosci 26:4095–4103.

Park S, Kramer EE, Mercaldo V, Rashid AJ, Insel N, Frankland PW, Josselyn SA (2016) Neuronal Allocation to a Hippocampal Engram. Neuropsychopharmacology 41:2987–2993.

Pavlides C, Greenstein YJ, Grudman M, Winson J (1988) Long-term potentiation in the dentate gyrus is induced preferentially on the positive phase of theta-rhythm. Brain Res 439:383–387.

Pedroni A, Minh do D, Mallamaci A, Cherubini E (2014) Electrophysiological characterization of granule cells in the dentate gyrus immediately after birth. Front Cell Neurosci 8:44.

Pignatelli M, Ryan TJ, Roy DS, Lovett C, Smith LM, Muralidhar S, Tonegawa S (2019) Engram Cell Excitability State Determines the Efficacy of Memory Retrieval. Neuron 101:274–284 e275.

Poirazi P, Brannon T, Mel BW (2003) Pyramidal neuron as two-layer neural network. Neuron 37:989–999.

Rathour RK, Narayanan R (2019) Degeneracy in hippocampal physiology and plasticity. Hippocampus 29:980–1022.

Santhakumar V, Aradi I, Soltesz I (2005) Role of mossy fiber sprouting and mossy cell loss in hyperexcitability: a network model of the dentate gyrus incorporating cell types and axonal topography. Journal of neurophysiology 93:437–453.

Schmidt-Hieber C, Jonas P, Bischofberger J (2004) Enhanced synaptic plasticity in newly generated granule cells of the adult hippocampus. Nature 429:184–187.

Schmidt-Hieber C, Jonas P, Bischofberger J (2007) Subthreshold dendritic signal processing and coincidence detection in dentate gyrus granule cells. J Neurosci 27:8430–8441.

Shors TJ, Dryver E (1994) Effect of stress and long-term potentiation (LTP) on subsequent LTP and the theta burst response in the dentate gyrus. Brain Res 666:232–238.

Shouval HZ, Bear MF, Cooper LN (2002) A unified model of NMDA receptor-dependent bidirectional synaptic plasticity. Proc Natl Acad Sci U S A 99:10831–10836.

Sjostrom PJ, Hausser M (2006) A cooperative switch determines the sign of synaptic plasticity in distal dendrites of neocortical pyramidal neurons. Neuron 51:227–238.

Sjostrom PJ, Rancz EA, Roth A, Hausser M (2008) Dendritic excitability and synaptic plasticity. Physiol Rev 88:769–840.

Srikanth S, Narayanan R (2015) Variability in State-Dependent Plasticity of Intrinsic Properties during Cell-Autonomous Self-Regulation of Calcium Homeostasis in Hippocampal Model Neurons. eNeuro 2:ENEURO.0053-0015.2015.

Wang Y, Rowan MJ, Anwyl R (1997) Induction of LTD in the dentate gyrus in vitro is NMDA receptor independent, but dependent on Ca2+ influx via low-voltage-activated Ca2+ channels and release of Ca2+ from intracellular stores. Journal of neurophysiology 77:812–825.

Yiu AP, Mercaldo V, Yan C, Richards B, Rashid AJ, Hsiang HL, Pressey J, Mahadevan V, Tran MM, Kushner SA, Woodin MA, Frankland PW, Josselyn SA (2014) Neurons are recruited to a memory trace based on relative neuronal excitability immediately before training. Neuron 83:722–735.

